# BRAWNIN: A sORF-encoded Peptide Essential for Vertebrate Mitochondrial Complex III Assembly

**DOI:** 10.1101/2020.01.31.926402

**Authors:** Shan Zhang, Chao Liang, Camille Mary, Baptiste Kerouanton, Joel Francisco, Narendra Suhas Jagannathan, Volodimir Olexiouk, Jih Hou Peh, Claire Tang, Gio Fidelito, Srikanth Nama, Ruey-Kuang Cheng, Caroline Lei Wee, Loo Chien Wang, Paula Duek Roggli, Prabha Sampath, Lydie Lane, Enrico Petretto, Radoslaw Sobota, Suresh Jesuthasan, Lei Sun, Lisa Tucker-Kellogg, Bruno Reversade, Gerben Menschaert, David A. Stroud, Lena Ho

## Abstract

The emergence of small open reading frame (sORF)-encoded peptides (SEPs) is rapidly expanding the known proteome at the lower end of the size distribution^1,2^. Here, we show that the mitochondria proteome is enriched for proteins smaller than 100 a.a. (defined as SEPs). Using a mitochondrial prediction and validation pipeline for small open-reading-frame (sORF)-encoded peptides (SEPs), we report the discovery of 16 endogenous mitochondrial SEPs (mito-SEPs) associated with oxidative phosphorylation (OXPHOS). Through functional prediction, proteomics, metabolomics and metabolic flux modeling, we demonstrate that BRAWNIN (BR), a 71 amino acid peptide encoded by the *C12orf73* gene, is essential for respiratory chain complex III (CIII) assembly. In human cells, BR is induced by the energy-sensing AMPK pathway, and its depletion impairs mitochondrial ATP production. *In vivo*, BR is enriched in muscle tissues and its maternal zygotic deletion in zebrafish causes complete CIII loss, resulting in severe growth retardation, lactic acidosis and early death. Our findings demonstrate that BR is essential for oxidative phosphorylation across vertebrate species. We propose that mito-SEPs are an untapped resource for essential regulators of oxidative metabolism.

## Main Text

SEP discovery is an emerging field, with recent SEPs implicated in diverse functions including cardiovascular and placental development, mTORC1 activation, calcium signaling, mRNA de-capping, membrane fusion and oocyte fertilization^3–8^. However, whether SEPs have programmatic functions enabled by their small size remains an open question. To this point, we observed that the mitochondria proteome, particularly at the inner mitochondrial membrane (IMM) (catalogued in Mitocarta 2.0^9^) is enriched for proteins smaller than 100 a.a. i.e. SEPs (p = 2.2×10^−16^) (Fig. 1a,b), and speculated that mitochondria are a hotspot for SEP function. To uncover novel mitochondrial-targeted SEPs (mito-SEPs) from the human SEP peptidome^10^, 2343 high quality human SEPs encoded by annotated short coding sequence (CDS) genes, long non-coding RNAs (lncRNAs), upstream ORFs (uORFs) and intergenic ORFs were selected for test of mitochondrial localization (Fig. 1c, details in SI). We first employed a three-prong strategy that identified a candidate mito-SEP on the basis of a mitochondrial gene expression signature (Fig. S1a); a mitochondrial targeting protein domain (Fig. S1b) or empirical detection of the SEP in mass spectrometric data of purified mitochondria (details in SI). Altogether, our mito-SEP discovery pipeline identified 173 candidates (Fig. 1c and Table S1), including 2 positive controls *MKKS uORF1* and *uORF2*^11^. HA-tagged mito-SEP candidates were experimentally validated by immunofluorescence of transfected HeLa cells (Fig. 1d). In total, we successfully validated protein expression for 88/173 sORFs (Table S1), defined as the ability of the ORF to generate a peptide with sufficient stability to enable detection (Fig. S1c-e). Of these, 23% (20/88) of successfully expressed peptides displayed unambiguous mitochondrial localization including the MKKS uORFs (Fig. 1d). Among the positive candidates were MTLN/MOXI and MIEF1 uORF which were subsequently independently discovered to have critical roles in mitochondria metabolism and mtDNA translation^12–14^, providing proof for the robustness of our pipeline. All in all, our screen identified 16 *bona fide* new mito-SEPs of unknown function (MSUF) encoded by uORFs, lincRNAs and short CDSs (Fig. 1d).

**Figure 1.**
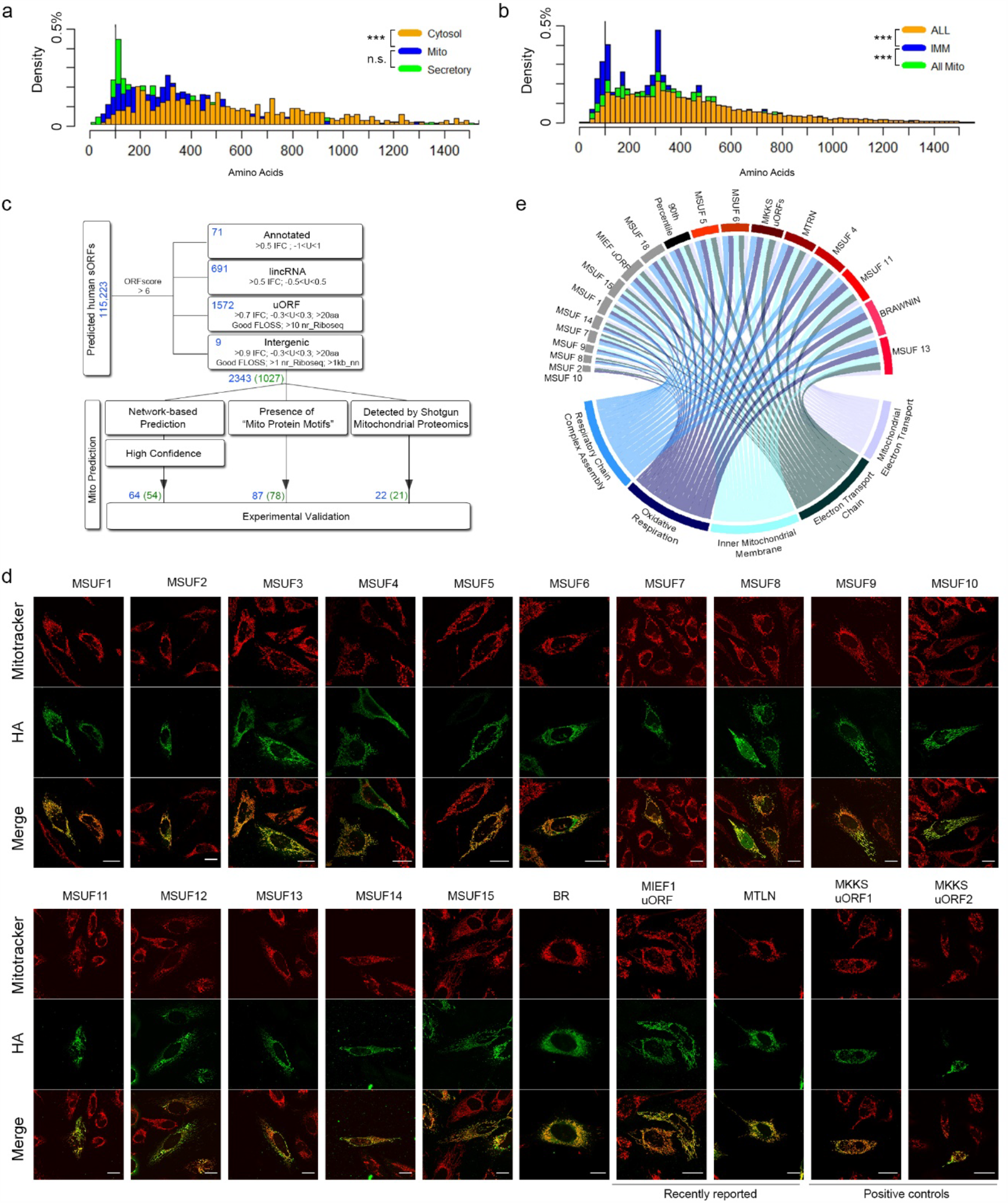
Mito-SEP prediction pipeline identifies novel endogenous mitochondrial SEPs. a. Size distribution of known proteins from Mitocarta 2.0 (Mito) and Uniprot proteins annotated as cytosolic or secretory. p-value by Mann-Whitney U-test. n.s. = not significant. b. Size distribution of Uniprot proteins (All) annotated to reside in mitochondria (All Mito) or inner mitochondrial membrane (IMM). p-value by Mann-Whitney U-test. c. Human SEP selection and mitochondrial prediction workflow. Blue numbers indicate number of SEPs and green numbers indicate number of genes encoding these SEPs. d. HA immunofluorescence (green) of mito-SEP candidates showing colocalization with Mitotracker Red (red) in HeLa cells. Novel SEPs are labeled as Mitochondrial SEP with Unknown Function (MSUF). Scale = 20 μm. e. Circular plot of expression correlation between Mito-SEPs and gene ontology (GO) gene sets that relate to oxidative phosphorylation.

Mitochondria are the principal sites of energy conversion where ATP is generated through the process of oxidative phosphorylation (OXPHOS). Notably, 63.5% (40/63) of known mitochondrial low molecular weight (MW) proteins function as assembly factors or core subunits of respiratory chain (RC) complexes that carry out OXPHOS (Fig. S1f). RC complexes are vice versa enriched for low MW proteins (Fig. S1g). Defects in OXPHOS are responsible for many inherited mitochondrial diseases and manifest in aging and metabolic disorders such as diabetes, heart disease, and cancer^15^. Of the 16 new mito-SEPs, the expression of 8 correlated with respiratory chain and electron transport function above the 90^th^ percentile (Fig. 1e), suggesting that half of them participate directly in oxidative phosphorylation.

We selected a mito-SEP encoded by the *C12orf73* gene, which we re-named *BRAWNIN (BR)* (Fig. 2a), for further characterization because its stable overexpression in U87MG significantly enhanced OXPHOS (Fig. S2a). The BR peptide is conserved in vertebrates (Fig. 2b) but not in yeast and nematodes. Although ribosome profiling detected 2 plausible BR peptides arising from alternative splicing (Fig. 2a and S2b), we confirmed that only the cDNA of the long isoform (P1) produced a stable 8 kD peptide, while the short isoform (P2) was remarkably less stable even when over-expressed (Fig. S2c). Endogenous BR was detected by western blotting with a custom α-BR antibody in HEK293T and HeLa cell lysates (Fig. 2c) and can be depleted by siRNAs targeting the *BR* transcript (Fig. 2d). Endogenous BR is enriched in mitochondrial-enriched fractions (Fig. 2e), and co-localized with Mitotracker in both HeLa and HEK293T (Fig. 2f-g, S2d). Altogether, these data confirm that BR is a nuclear-encoded peptide that is imported into the mitochondria. Consistent with this observation, BR can be detected in human and mouse skeletal muscle and human cardiac muscle (Fig. 2h-k), where it displays a staining pattern characteristic of the mitochondrial network. Moreover, BR protein abundance in mouse tissues correlates with mitochondrial content, being high in brown adipose, cardiac and skeletal muscle but virtually undetectable in white adipose tissue (Fig. S2e). Because mitochondria are highly compartmentalized, the molecular function of a protein determines its sub-mitochondrial location and vice versa^16^. We first established that BR is membrane-bound by sodium carbonate extraction (Fig. 2l). Next, using differential detergent extraction, we determined that BR resides in the IMM and not in the outer mitochondrial membrane (OMM) (Fig. 2m). Proteinase K protection assays further suggest that the C-terminus of BR faces the intermembrane space (IMS) (Fig. 2n). In contrast to its clear mitochondrial localization, human BR lacks a classical mitochondrial targeting sequence (MTS); its N-terminus is instead predicted to function as a secretory signal peptide by Signal-P^17^. Indeed, when overexpressed, BR can be secreted (Fig. S2f). However, we were unable to detect secretion of the endogenous protein from HEK293T cells grown under standard conditions (data not shown). Hence, the N-terminal “signal peptide”, which is not cleaved (Fig. S2g), functions as a mitochondrial targeting single-pass transmembrane domain (TMD) to anchor endogenous BR into the IMM. Consistent with this, BR-P2, which contains the full TMD of BR-P1, is similarly targeted to the mitochondria despite its relative instability (Fig. S2h). Removal of the predicted TMD from BR (BR^ΔN25^), while causing its near complete destabilization (Fig. S2i), targets BR^ΔN25^ to the cytosol instead of the mitochondria (Fig. S2j).

**Figure 2.**
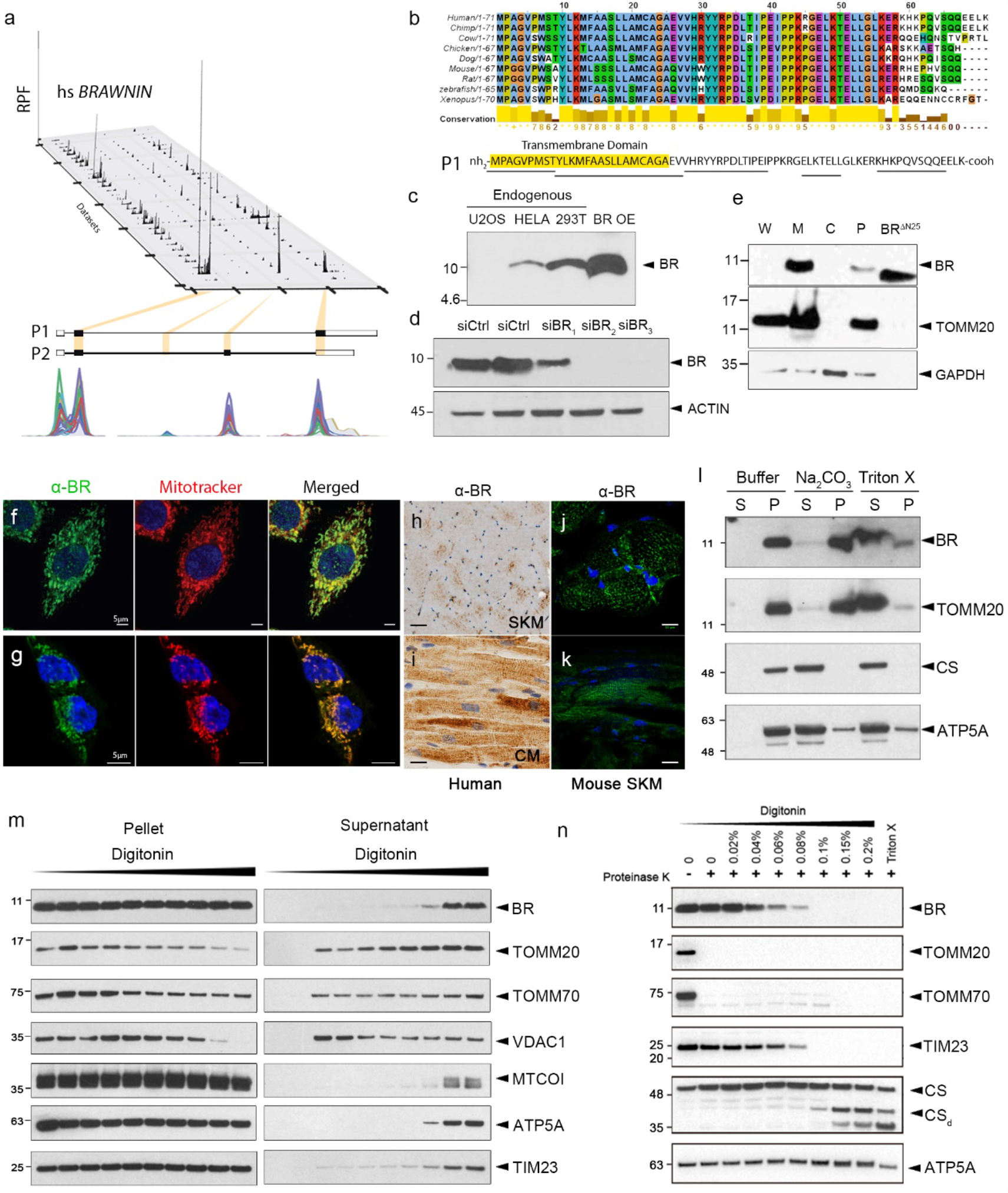
BRAWNIN (BR) is a conserved SEP at the inner mitochondrial membrane. a. Ribosome-sequencing (Ribo-seq) reads from the *BRAWNIN* gene *(C12orf73)* across 31 human cell lines. P1 and P2 refer to the 2 potential ORF isoforms generated by alternative splicing. b. BR (P1) peptide is conserved throughout vertebrate evolution and possesses an N-terminal hydrophobic region that is predicted to be a transmembrane domain or signal peptide. Underlined regions are peptides detected by mass spec. c. Endogenous BR is detected in HeLa and HEK293T with an α-BR antibody. OE=overexpression d. BR knock down using siRNAs in HEK293T. e. Mitochondrial fractionation of HEK293T. W: whole cell; M: mitochondria enriched fraction; C: cytosol; P: nuclear fraction and debris, including un-extracted mitochondria; sBR: synthetic BR peptide. f. Endogenous BR detected by α-BR in HeLa costained with mitotracker. Scale = 5 μm. g. Endogenous BR detected by α-BR in HEK293T costained with mitotracker. Scale = 5 μm. h. α-BR immunohistochemistry (IHC) in human SKM, cross section. Scale = 20 μm. i. α-BR IHC in human cardiac muscle (CM), longitudinal section. Scale = 20 μm. j. α-BR IF in mouse SKM, cross section. Scale = 12 μm. k. α-BR IF in mouse SKM, longitudinal section. Scale = 12 μm. i. Extraction of endogenous BR from HEK293T mitochondria with buffer alone, Na2CO3 or Triton X. S=soluble; P= pellet. m. Extraction of endogenous BR from HEK293T mitochondria using increasing concentrations of digitonin. TOMM20, TOMM70 and VDAC1 are OMM proteins, are MTCO1, ATP5A and TIM23 are IMM associated. n. Proteinase K protection assay of HEK293T mitochondria. The α-BR epitope is C-terminal to the predicted transmembrane domain. TOMM20 and TOMM70 have cytosolic domains. TIM23 has an IMS domains while both CS and ATP5A are in the matrix.

Proteins in the IMM participate in a wide variety of processes, most commonly OXPHOS, calcium import, and the transport of proteins and metabolites. Co-expression analysis by weighted gene co-expression network analysis (WGCNA)^18^ supports the notion that BR participates in electron transport and OXPHOS in multiple human organs and species (Fig. 3a, S3a, Table S2). Components of the respiratory chain are dynamically regulated by energy status. In response to a reduction of cellular ATP:ADP ratio, such as during nutrient depletion or exercise, the energy-sensing AMP-activated protein kinase (AMPK) pathway restores energetic balance by stimulating ATP-producing catabolic pathways and inhibiting energy-consuming anabolic processes^19^. As such, we hypothesized that BR levels would be regulated by cellular energy status. Indeed, activation of AMPK using 5-aminoimidizole-4-carboxamide-1-β-D-riboside (AICAR)^20^ robustly increased protein levels of BR (Fig. 3b). Likewise, BR increased in response to glucose, serum and fatty acid starvation, in parallel with AMPK activation marked by phosphorylation of Acetyl-CoA carboxylase (ACC), a direct AMPK target^21^ (Fig. 3c). In mouse myotubes, enforced expression of PGC-1α, the AMPK-activated regulator of mitochondrial biogenesis^22^, robustly induced *Br* expression (Fig. S3b). These data together indicate that BR is under the regulatory control of the AMPK-PGC-1αenergy homeostasis axis, and predict that a loss of BR would impair cellular bioenergetics and mitochondrial ATP production. To investigate this, we silenced *BR* in U87MG glioblastoma cells (a highly oxidative cell type) using both transient siRNA and stable shRNA-mediated approaches. Indeed, BR depletion significantly decreased basal and maximal respiration, spare capacity and ATP production in both si*BR* and sh*BR* U87MG (Fig. 3d-f, S3c). Likewise, HEK293T grown in galactose displayed the same reduction in basal respiration (Fig. S3d), which sufficiently perturbed cellular ATP:ADP ratio to activate AMPK (Fig. S3e). These data confirm that *BR* is essential for cellular bioenergetics, potentially through direct regulation of the electron transport chain.

**Figure 3.**
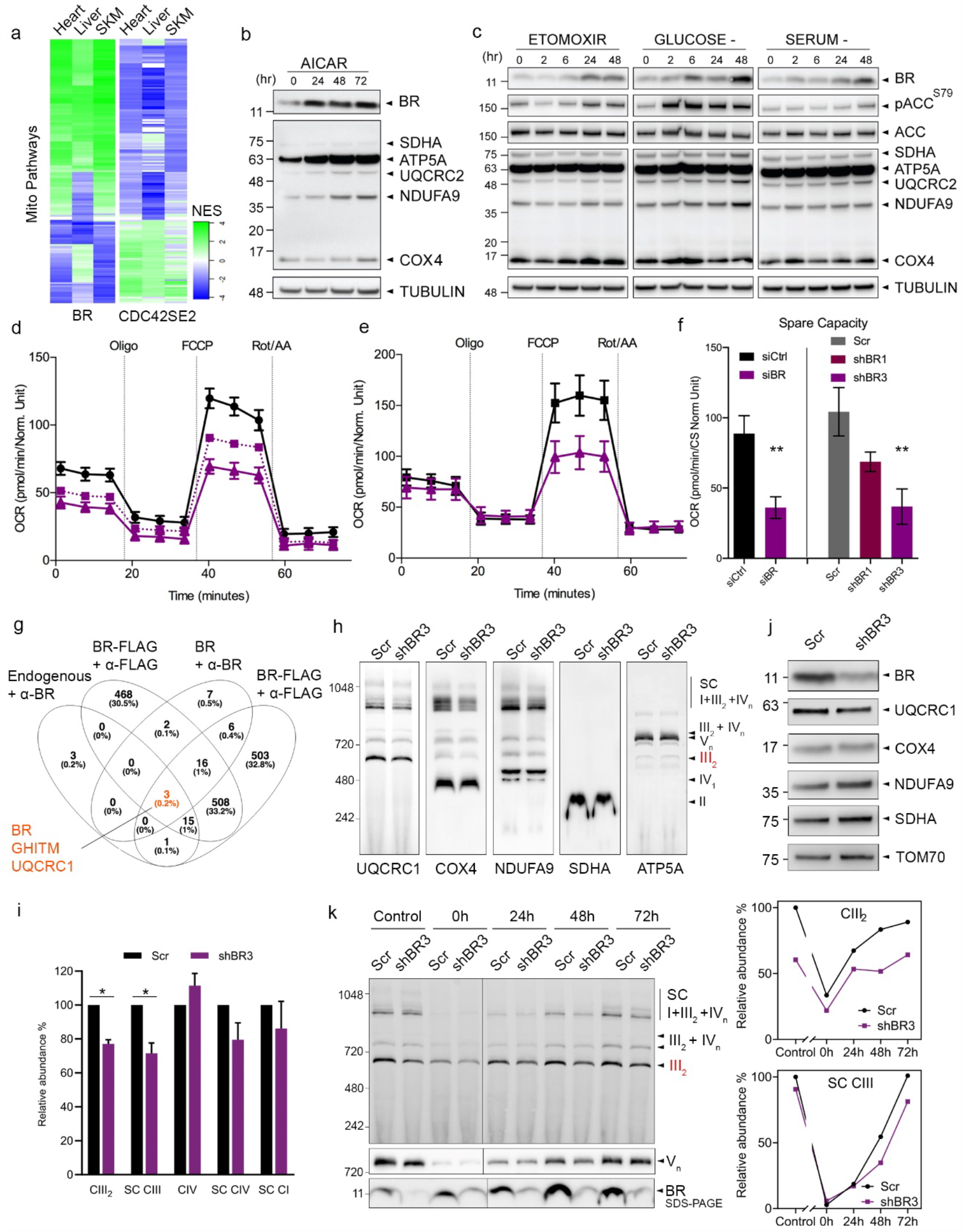
BR is an AMPK target that potentiates OXPHOS and respiratory complex assembly. a. Heatmap depicting GSEA NES scores of mitochondrial gene sets (see Table S2) in the ranked list of pairwise correlations between genes in the human liver, heart or skeletal muscle (SKM) with either *BR* gene or *CDC42SE2*, a representative non-mitochondrial candidate SEP from Fig. 1D. b. SDS-PAGE of HEK293T treated with AMPK activator AICAR (500 μM) for the indicated time periods. c. SDS-PAGE of HEK293T treated with indicated starvation regimes for the indicated time periods. ETOMOXIR was used at 3μM. d. Seahorse mito stress test analysis of control versus stable *BR* knockdown U87MG cells. Oxygen consumption rate (OCR) reads were normalized with citric synthase (CS) activity. Data are mean and SEM of 6 technical replicates. Experiment is representative of 2 biological replicates. Oligo, oligomycin; FCCP, carbonyl cyanide 4-(trifluoromethoxy)phenylhydrazone; Rot/AA, rotenone/antimycin A. e. Analogous to d. except using short-term (72 hour) siRNA-mediated depletion of BR. Experiment is representative of 3 biological replicates each with 6 technical replicates. f. Spare respiratory capacity calculated from experiment in d. and e. Data are mean and SEM of 2 and 3 biological replicates for shRNA and siRNA-mediated depletion, respectively, each with 6 technical replicates. p-values from unpaired t-test (α=0.05). g. Venn diagram depicting intersections of proteins identified in 4 co-IP/mass spec experiments of BR in 1% DDM-solubilized HEK293T mitochondria. Endogenous BR was immunoprecipitated with α-BR, overexpressed BR-FLAG with α-FLAG, overexpressed BR untagged with α-BR. h. BN-PAGE of mitochondria from U87MG stably transduced with scramble control (Scr) or shRNA targeting *BR* (shBR3). UQCRC1 densitometric signal in the SC range was normalized to ATP5A to reveal a 30% reduction in shBR3 SC compared to Scr SC. l. Relative abundance of CIII_2_, supercomplex assembling CIII (SC CIII), CIV, supercomplex assembling CIV (SC CIV), supercomplex assembling CI (SC CI) in scramble control (Scr) or shRNA targeting *BR* (shBR3) transduced U87MG cells. Error bars indicate SEM of three independent replicates. j. SDS-PAGE of mitochondria from control and *sh*BR3 U87MG. k. BN-PAGE of mitochondria from U87MG cells after 8 days of doxycycline treatment at 15 μg/ml. Mitochondria were harvested at indicated times after removing doxycycline from the culture media. Protein abundances of BR in corresponding samples were determined by SDS-PAGE.

To deduce the molecular function of BR, we constructed its interactome in HEK293T mitochondria by co-immunoprecipitating either endogenous or overexpressed BR followed by mass spectrometry (IP/MS). At the intersection of 4 independent IP/MS experiments were only 3 proteins: BR, GHITM, and UQCRC1 (Fig. 3g, Table S3). UQCRC1 is a core component of respiratory chain CIII^23^ while GHITM has been proposed to regulate cytochrome c release during apoptosis^24^. The interaction with GHITM could not be validated with several commercially available antibodies (data not shown). In contrast, the interaction between BR and UQCRC1 detected in human cells was validated by co-IP/western blot in digitonin-solubilized mouse heart mitochondria expressing a Br-FLAG construct (Fig. S3f). These data suggest that BR, an IMM peptide, directly interacts with the respiratory chain at CIII. Among all genes encoding RC proteins, *BR* transcript expression correlates most strongly with those encoding CIII assembly factors (Table S4). On this basis, we hypothesized that the observed respiratory defects in si/sh*BR* U87MG are caused by CIII deficiency. To test this, we turned to biochemical analyses using blue-native (BN) PAGE, which separates RC complexes into their free forms and forms residing in higher order supercomplexes (SCs or respirasomes)^25^. Indeed, in *shBR* U87MG, the levels of fully assembled CIII_2_ dimers and SCs were reduced by 25% at steady state (Fig. 3h,i). Total UQCRC1 protein levels were similarly decreased, while core components of other RC complexes were unaffected (Fig. 3j). As an assembly factor, we predicted that BR deficiency would impair the rate of CIII biogenesis during a mitochondrial challenge. We therefore assessed the assembly kinetics of CIII_2_ and CIII-containing SCs following 8 days of doxycycline-induced reversible inhibition of mitochondrial translation. Indeed, BR deficiency delayed the recovery of CIII_2_ levels and SCs after removal of doxycycline (Fig. 3k), demonstrating its requirement as a CIII assembly factor. Altogether, our results indicate that human BR responds to energy status by promoting CIII biogenesis to enhance bioenergetic efficiency for maximal ATP production.

*BR* is highly conserved in vertebrates, suggesting that its role in maintaining CIII integrity is fundamental and required throughout evolution. To understand the *in vivo* requirement of BR in animal physiology, we turned to the zebrafish where *br* is most highly expressed in skeletal and cardiac muscle (Fig. 4a). Using Crispr/Cas9, we generated *br* knockout animals (KO) by removing the entire protein-coding ORF contained in exons 2 and 3 (Fig. 4b) which resulted in a complete protein null (Fig. 4c). *br* KO larvae were produced at Mendelian ratios (Fig. 4d) with no gross development defects within the first 5 days besides a delay in swim bladder inflation (Fig. S4a,b). Muscle formation was normal with no overt signs of myopathy or dystrophy (Fig. S4c). However, if left unsegregated, zygotic *br* KOs (*z*KO) cannot be recovered by 65 days post fertilization (dpf) (Fig. 4d). We segregated larvae from intercrosses at 3 dpf and allowed them to grow at low density. Under this regime, *z*KO juveniles survived into adulthood, but displayed growth retardation from 28 dpf (Fig. 4e) that resulted in a large difference in body size by 65 dpf (Fig. 4f). *z*KOs are sub-fertile, and can generate maternal zygotic knockout embryos (*mz*KO) albeit at a very low frequency (about 1/20 mating pairs). Because Br is a mitochondrial peptide, it is maternally deposited in oocytes. *mz*KOs therefore lack any residual Br protein that is available to *z*KOs during the initial phases of larvagenesis. Consequently, *mz*KO had a more severe phenotype compared to *z*KOs (Fig. 4e). By 6 dpf, *mz*KO larvae were visibly less active despite fully inflated swim bladders (Fig. S4d). By 7 dpf, at the onset of feeding, they ate significantly less than WT larvae (Fig. S4e). By 11 dpf, they displayed a dramatic reduction of directional swimming and distance swam compared to WTs (Fig. 4g and S4f). Consequentially, *mz*KO larvae do not increase in body length even under low density segregated growth conditions, and do not survive past 42 dpf (Fig. 4e). To test if the observed physiological defects are due to CIII deficiency, we employed an independent approach of profiling the metabolomic perturbations in whole *mz*KO animals at 5 dpf. Targeted metabolomics analysis revealed a 2-fold increase in the levels of lactate in *mz*KO larvae (Fig. 4h) and a 3-fold increase in the citric acid cycle intermediate succinate as compared to WT larvae (Fig. 4i). Lactate accumulation is caused by decreased pyruvate oxidation secondary to mitochondrial deficiency and is frequently used as a diagnostic symptom for patients with mitochondrial diseases^26^. To understand how these perturbations could be related to CIII deficiency, we turned to MitoCore, a metabolic flux model that simulates over 300 mitochondrial reactions in a human cardiomyocyte using flux balance analysis^27^. By blocking each reaction in turn, and simulating 5000 feasible flux states for each possible blockage, we found that inhibiting CIII and CIV (but not CII) flux caused the largest, most biologically plausible increase in succinate and lactate (as inferred from the export rate of the two metabolites) (Fig. 4j, S4g-i). Similarly, when we deleted each of the 371 genes in the MitoCore model, the deletion of genes encoding CIII and CIV components caused the largest increase in succinate and lactate (Table S5). Since CIII flux is the only upstream determinant of CIV flux in MitoCore, the observed CIV defects are likely driven by CIII defects. Strikingly, BN-PAGE analysis of *br* KO mitochondria revealed a 94 % reduction of assembled CIII complexes as measured by Uqcrc1 densitometry (Fig. 4k), while total Uqcrc1 was reduced by 77 % (Fig. 4l). These data provide confirmation for the *in vivo* requirement of Br in CIII assembly and/or stability seen in human U87MG cells. Furthermore, spectrophotometric measurement of RC enzymatic activities^28^ revealed that all CIII-dependent activities were indeed impaired in *z*KO muscle homogenates (Fig. S4j). Intriguingly, CIV activity, as measured by uncoupled electron flow in response to TMPD/ascorbate, was reduced by 5-fold in *br* KO mitochondria (Fig. 4m). However, since CIV protein levels (Mtco1 and Cox4) are not affected (Fig. 3h, 4l-m), CIV impairment is unlikely to be the primary defect in *br* KO animals. Rather, reduced CIV activity is likely caused by the near complete loss of CIII_2_ + IV super-assemblies^29^ secondary to the loss of CIII (Fig 4k). Consistent with a direct role of Br as a CIII assembly factor *in vivo*, BN-SDS PAGE demonstrated co-migration of endogenous Br with CIII in zebrafish mitochondria (Fig. S4k,l). The functional consequences of the observed molecular deficiencies were dramatic. By performing Seahorse respirometry on isolated mitochondria, we found that OXPHOS and ATP production in *br* KO mitochondria were reduced to half that of WT mitochondria (Fig. 4n). At the physiological level, this impairment significantly reduced whole animal respiration *br* mz KO compared to WT larvae even as early as 1 dpf (Fig. 4o), leading to a failure to thrive and early death (Fig. 4e). Altogether, our data demonstrate that the loss of *Br* in zebrafish causes overt and lethal mitochondrial deficiency due to the loss of CIII in the respiratory chain.

**Figure 4.**
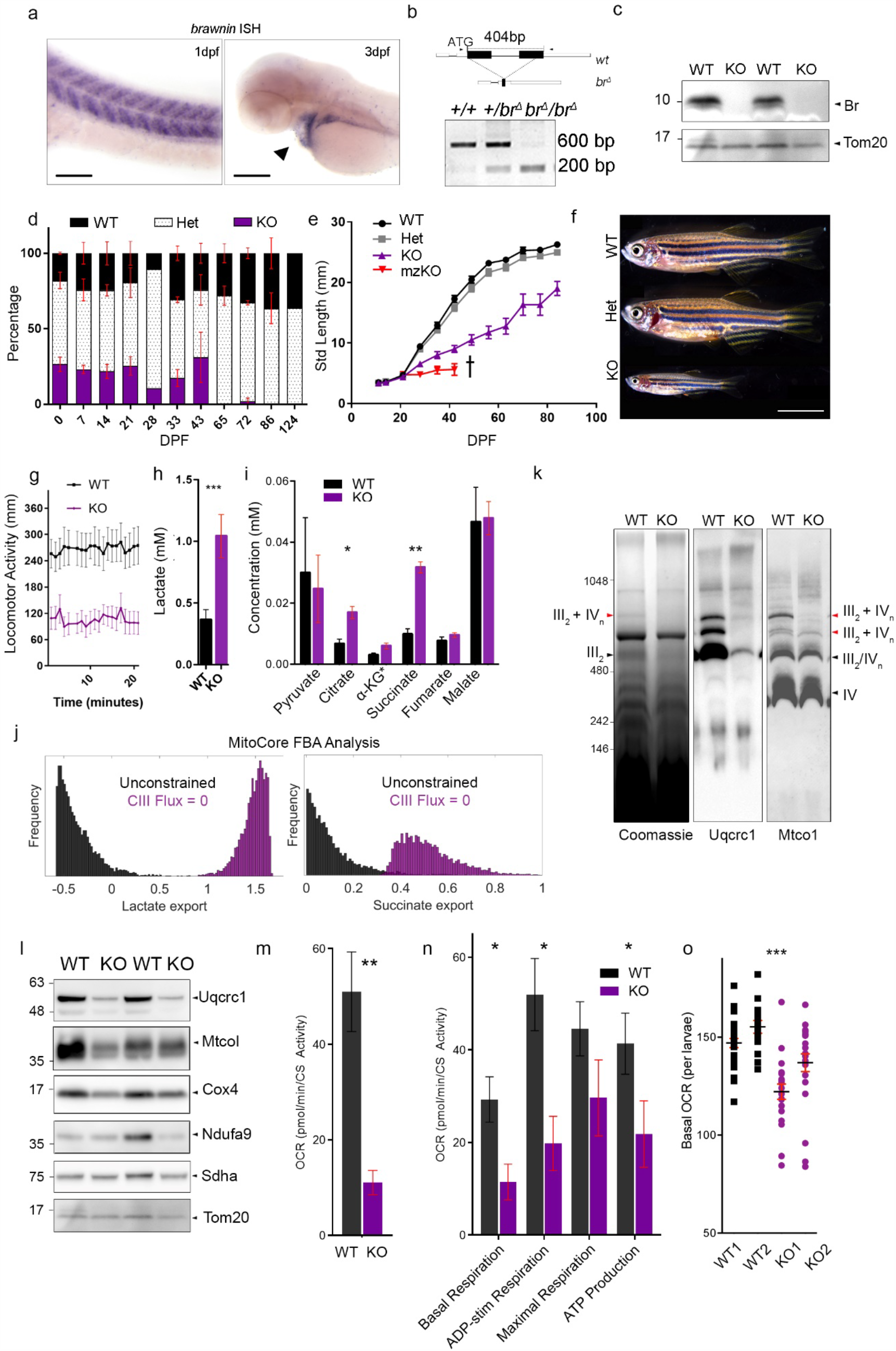
Knockout of *Br* in zebrafish causes lethal mitochondrial deficiency. a. *In-situ* hybridization (ISH) of *brawnin (br)* in zebrafish larvae. dpf = days post fertilization. Left panel: scale = 100 μm, right panel: scale 200 μm. b. Deletion strategy of *zf br* by Crispr/Cas9 (top); PCR of genomic locus with indicated primers spanning deletion site. c. Western blot analysis of purified mitochondria from adult WT and KO (homozygous *br*^Δ^/*br*^Δ^) skeletal muscle using zebrafish-specific α-Br. d. Mendelian percentages of offspring from heterozygous (het) intercrosses genotyped at the indicated dpf. Data represent the mean of at least 2 clutches per time point (except 28 and 124 dpf) with at least 15 animals per clutch. Error bars indicate SEM. e. Standard length (mm) measurements of WT, Het and zygotic KO larvae generated from an F_2_ intercross genotyped and segregated at 3 dpf. n > 20 per genotype at the beginning of the experiment. Maternal zygotic KOs (mzKOs) were generated from a homozygous incross and measured until 42 dpf when all animals succumbed to disease. Data are representative of 4 independent experiments from 2 independent Cas9/gRNA injected *br*^Δ^ KO allele founders. Error bars indicate SEM. f. Representative images of WT, Het and zygotic KO clutch-mates at 65 dpf. Scale = 5 mm. g. Basal motility monitoring continuous swim tracking of WT and KO larvae at 11 dpf. h. Lactate concentrations in 5 dpf WT and mzKO larvae. Data represent mean and SEM of 4 independent clutches each with 50 larvae per genotype. p-values from paired t-test (α=0.05). i. Citric acid cycle organic acid concentrations in 5 dpf WT and mzKO larvae. Data represent mean and SEM of 4 independent clutches each with 50 larvae per genotype. p-values from paired t-test (α=0.05) with Bonferroni correction. *α-KG concentrations are a 10-fold multiple to enable visualization on the same scale. j. Distributions of the export rate of lactate and succinate (5000 flux samples) comparing the unconstrained MitoCore model versus the CIII-blocked model derived by MitoCore flux balance analysis (FBA). k. BN-PAGE and western blot analysis of matched WT and zKO adult skeletal muscle (SKM) mitochondria. Position of free III_2_ (black) and III_2_ + IVn (red) supercomplexes are indicated by arrowheads. l. SDS-PAGE of purified WT and zKO skeletal muscle mitochondria from (L). The Tom20 blot is repeated from Figure 4C as they are from the same experiment. m. Uncoupled electron flow measured by Seahorse of purified mitochondria from WT and zKO skm using N,N,N’,N’-Tetramethyl-p-Phenylenediamine (TMPD)/Ascorbate as the electron donor, read out as CS-normalized OCR. Data represent mean and SEM of 4 biological replicates. p-values from paired t-test (α=0.05). n. Coupled respiratory analysis of isolated SKM mitochondria in the presence of succinate + rotenone, read out as CS-normalized OCR. Data represent mean and SEM of 4 biological replicates. p-values from paired t-test (α=0.05). o. Basal respiratory rate of free moving 1 dpf WT and *mz*KO larvae. Each dot represents one animal. Data represent mean and SEM, p-values from unpaired t-test (α=0.05).

The mechanism with which BR promotes CIII assembly calls for greater in-depth investigation. The assembly of Complex III in mammals is poorly understood and largely inferred from studies performed in yeast^30^. In zebrafish, loss of Br caused complete attenuation of the fully assembled 500 kDa CIII dimer (i.e. III_2_), with no evidence of immature pre-CIII_2_ assemblies seen in mutants of late assembly factors^31–33^, arguing that Br acts early in the biogenesis and known sequence of CIII assembly before dimerization of assembly intermediates^31^. BR is highly responsive to energy status. Electrons transferred from the oxidation of divergent fuel types converge at CIII. Because CIII is the center of the respiratory chain, we speculate that BR, acting as an early CIII assembly factor, coordinates the bioenergetic output of the respiratory chain with nutrient availability to maintain energy homeostasis. Recently, CIII integrity has been implicated in regulatory T cell function and the prevention of autoimmunity^34^, raising the intriguing possibility that BR can also coordinate nutrient availability with immune function. Lastly, congenital CIII defects resulting from mutations in CIII subunits or assembly factors cause very rare but severe mitochondrial disorders such as GRACILE syndrome^35,36^, characterized by growth retardation, lactic acidosis, aminoaciduria and early death. The phenotypic similarities of the *br* KO zebrafish set the basis to screen unsolved cases of mitochondrial diseases like GRACILE syndrome for pathogenic variations in the *BR* (*C12orf73*) gene.

In conclusion, by performing a screen of the SEP peptidome, we report the discovery of BRAWNIN, a novel mitochondrial peptide that is central to vertebrate oxidative phosphorylation. This emerging theme of the preponderance of SEPs in the mitochondria is seen also in the human heart^37^, pointing to the *in vivo* significance of mito-SEPs in human physiology. Of high priority is deducing the contribution of the other 15 mito-SEPs to oxidative metabolism. Our work demonstrates the promise and value of mining and characterizing the sORF-encoded peptidome, which should motivate similar efforts to functionally characterize remaining novel SEPS, also known as <u>S</u>mall G<u>E</u>ms hidden in <u>P</u>lain view.

## Methods

### sORF Selection Criteria

sORF-encoded peptides (SEPs) were filtered using ribosome profiling-based metrics derived from sORFs.org, adapting the guidelines from Olexiouk et. al.^38^. 2343 human SEPs encoded by annotated short CDS genes, lincRNAs, upstream ORFs (uORFs) and intergenic ORFs that passed ribosome sequencing quality thresholds were selected for mitochondrial prediction (Ensembl annotation bundle v92 is applied). From the pool of 115,223 predicted sORFs, a minimum ORF-score of 6 ^39^ was imposed on nucleus encoded sORFs. Extra filtering criteria are set separately dependent on the sORF annotation. For the sORFs annotated in CDS genes, a minimum threshold of 50% in-frame coverage and a coverage uniformity between +1 and −1 was imposed, whereas for sORFs located on lincRNA, an in-frame coverage of 50% and a coverage uniformity between −0.5 and 0.5 was imposed. For sORFs located in the 5’-UTR region of protein coding genes, a minimum in-frame coverage of 0.7, a coverage uniformity between −0.3 and 0.3, a minimum length of 20 a.a. and a good FLOSS categorization was imposed ^40^. Also, these sORFs should be identified/predicted in at least 10 different RIBO-seq datasets. For intergenic sORFs (i.e. in between genes), a minimal up- and down-stream gene distance of 1000 nucleotides (>1 kb_nn), a minimal in-frame coverage of 0.9, a coverage uniformity between −0.3 and 0.3, a minimum length of 20 a.a., a good FLOSS categorization, and a minimum number of identifications in at least 1 RIBOseq dataset was imposed. This resulted in a pool of 2343 sORFs encoded from 1027 different genes (Fig. 1C). See Supplemetary text for more explanation of selection parameters.

### Mitochondrial Prediction by Weighted Correlation Network Analysis (WGCNA) and Gene Set Enrichment Analysis (GSEA)

RNAseq datasets from 3 human tissues known to be enriched in mitochondrial activity were selected to enable mitochondrial functional prediction: non-neoplastic liver (21 samples from GSE94660 – RPKM normalized), normal heart left ventricular tissue (92 samples - VST counts)^41^ and normal skeletal muscle (84 samples from GSE120862 – DESeq normalized counts). To compare those mitochondria-enriched tissues with a non-enriched one, a sun-protected skin dataset (91 samples from GSE85861 – Normalized counts) was selected. To clean the datasets, genes with at least one read in more than half of the samples were kept except for the heart dataset. The healthy heart dataset is coupled to a dilated cardiomyopathy (DCM) dataset and genes with at least one read in more than 5% of the samples were kept. To distinguish between a mitochondrial and a non-mitochondrial gene, we selected 1158 *bona fide* mitochondrial genes (“mito”) (from Human MitoCarta2.0) for comparison against 957 nuclear transcription factors (TF) randomly selected from www.tfcheckpoint.org (“non-mito”) ^9,42^. These form the “classifier genes”. We further selected 56 known MitoCarta genes that encoded proteins < 100 a.a. (i.e. mito-SEPs) and 83 annotated nuclear and secreted proteins <100 a.a. (i.e. non-mito SEPs) which formed the mito vs non-mito SEP. These form the “training genes”. We performed WCGNA with the WGCNA package in R ^18,43^ for all classifier and training set genes in all 3 datasets. i.e. we ranked all genes in a dataset according to their Spearman’s correlation with each test gene. Next, we performed GSEA ^44^ using 7 gene set collections from MSig DB (Broad Institute) ^45,46^: c2.cp.reactome, c2.cgp, c2.cp.kegg, c5.cc, c5.bp, c5.mf and h.all, which gather 10260 gene sets. Using fgsea package in R ^47^, the NES score of each gene set was calculated for each ranked list. NES was set to 0 if the GSEA output p-value was not significant (p>0.05). To decrease noise and increase specificity to mitochondrial activity, only gene sets with at least 100 mito genes with an absolute NES above 3 were kept and gene sets containing more than 500 genes were removed. On average, each gene had 63, 85 and 118 NES values in the heart, liver and SKM dataset respectively. These scores were used as variables for principal component analysis (PCA) using stats package. Altogether, each PCA analysis contained 3090 observations (1956 classifier + 139 training + 995 SEP candidate genes). Only 995 of the 1027 genes encoding 2343 SEP candidates were expressed in any one of the 3 datasets and could be tested by this method. Next, K-means clustering was applied to the PCA for a cluster number varying from 2 to 15. For each iteration, clusters were ranked from the highest percentage of mito genes to the lowest. To know if a cluster should be considered as positive or negative for mitochondrial identity, each cluster’s mito percentage was tested as a potential threshold. The threshold which led to the highest number of correct prediction of the training set was kept. Clusters with a mitochondrial percentage below the threshold were considered negative, otherwise positive. Candidates were given a binary score of 0 or 1 depending on whether they belonged to a negative or positive cluster respectively. This process of binary scoring was made for every cluster number, and each SEP candidate was given an average k-means score. K-mean score was set to −1 if the gene was not expressed in the dataset. Based on the k-means scores of the training set, it appeared that 75% was the minimum average k-means score needed for a peptide to be predicted as mitochondrial. To be more stringent on the selection we added the criteria of confidence level. “High confidence” meant that a SEP candidate required an average k-means score above 75% in all RNAseq where it is expressed. Refer to Supplemental Text for more information.

### Mitochondrial Prediction by Mitochondrial Targeting Motif Prediction

Mitochondrial targeting motifs were predicted in peptides that were not selected by WGCNA/ GSEA. Different types of mitochondrial targeting motifs were predicted using separate algorithms. Classical mitochondrial targeting sequences (MTSs) were captured by the consensus prediction of TargetP (mitochondria, Reliability class 1 & 2) and Mitofate (score>0.8) or Mitoprot (score>0.8) ^48–50^. Transmembrane domains (TMDs) with modest hydrophobicity also serve as mitochondrial targeting signal, but they are poorly predicted by algorithms used above because these sequences are not cleavable. These hydrophobic α-helices were predicted by TMHMM or SignalP (4.1 and 5) ^48,51^. To eliminate secretory proteins, the maximum TMD hydrophobicity was set at 3 ^52^, as we noticed that mitochondrial TMDs were on average less hydrophobic compared to non-mitochondrial TMDs ^53^. In addition, mitochondria intermembrane proteins often contain twin cysteine pairs (two C9XC, two C3XC, or one C9XC and one C10XC, where X refers to any residues in between two cysteine residues) ^54^, and these internal signals are generally missed by prediction algorithms. We thus included peptides with these characteristic cysteine pairs in the screen.

### Mitochondrial Prediction by Mitochondrial Peptide RESPIN

Quantitative mass spectrometry data on isolated mitochondria are available at the PRIDE public proteomics data repository (accession number PXD004666) ^55^ and downloaded using the PRIDE API ^56,57^. Subsequently, the raw data files were converted and filtered using MSconvert ^58^. Next, the SearchGui tool ^59^ was used to perform the peptide to spectrum matching, based on a combination of two database search algorithms X!tandem ^60^ and MSGF+ ^61^ against a custom sequence database. This database consists of (i) the UniProt human functional proteome including all isoforms ^62^, (ii) the human sORFs from the sORFs.org database ^63^, and (iii) the cRAP database (https://www.thegpm.org/crap/). Search engine parameters were adapted from the original study ^55^. Matched PSMs were validated using PeptideShaker ^64^, only PSM solely mapping to sORF peptide sequences with a spectrum coverage of at least 30% and FDR<0.01 were retained. After selecting peptides by the three strategies, redundant isoforms were manually removed if they share identical sequences with the longest isoform encoded by the same gene, with ribosomal periodicity information been considered.

### SEP Transient Over-expression

The open reading frames (ORFs) of selected candidates were synthesized and inserted into the NheI-XhoI site of a pcDNA3.1 (+) vector, under the control of the CMV promoter. Peptides were tagged with a single HA tag distal to their predicted targeting motifs or c-terminus if no motif was predicted. ORFs with non-AUG start codons were replaced with a AUG start codon to avoid the need for including cognate 3’UTRs required for recognizing near-cognate start sites ^65^. HeLa cells were seeded in 24-well plates and transfected with FuGENE (Promega) and stained as described in the immunofluorescence section.

### Zebrafish Br Knockout Generation

zf*Brawnin* is encoded by the si:ch211-68a17 gene which contains 3 exons with the ORF in exons 2 and 3. EST mining on NCBI and ENSEMBL mapping to this region in the zebrafish genome did not reveal the presence of a duplicated allele or paralogue. To completely excise the ORF, we used ZiFiT ^66^ to design two gRNAs 5’ and 3’ to the ORF with recognition sites underlined. ATG in bold marks the translation start site.

5’ zfBR_gRNA : 5’GAAATTAATACGACTCACTATA<u>GG***ATG***CCAGCAGGCGTATCT</u>GTTTTAGAGCTAGAAATAGCGTTTTAGAGCTAGAAATAGCAAGTTAAAATAAGGCTAGTCCGTTAT CAACTTGAAAAAGTGGCACCGAGTCGGTGCTTTT 3’

3’ zfBR_gRNA: 5’GAAATTAATACGACTCACTATA<u>GAACTGCGAACAGAACTTCT</u>GTTTTAGAGCT AGAAATAGCAAGTTAAAATAAGGCTAGTCCGTTATCAACTTGAAAAAGTGGCACCGAGTCGGTGCTTTT3’

gRNAs were generated from gBLOCKS with MEGAshortscript™ T7 Kit (Invitrogen, AM1354). Following DNaseI treatment and precipitation with NH_4_Ac and ethanol, gRNAs were resuspended in 40 μl pure water. 200 pg of each gRNA was injected along with 300 pg of purified recombinant Cas9 protein (a kind gift of Harwin Sidik) into 1 cell stage AB larvae. F0 founders were detected by genotyping with the following primers:

5’AAAATTTCTCATCCTTAGCATAACATC3’ (LHW4_zfBr_5’_F)
5’AGAACTTGCAGTTGGCATCA 3’ (LHW7_zfBr_3’_R)

WT undeleted band yields a 624 bp band while the deleted allele (*brΔ)* yields a 220 bp with HotStart Taq Polymerase (Qiagen). 2 *brΔ* founders (ΔBig and ΔSmall) with complete deletions were brought forward for characterization. Both were backcrossed to AB until F2 before phenotypic investigations began. All data showed in the paper are generated from F2 and F5 incrosses. Growth retardation phenotype is stable throughout generations and reproducible to F6 when this manuscript was prepared. Both founders showed similar growth defects. All data presented are derived from ΔBig but are representative of ΔSmall, which had a more severe phenotype.

### Zebrafish Husbandry and Growth Tracking

Zebrafishes were reared under standard conditions, 28°C with approved protocols following regulations stipulated by the IACUC committee of the National University of Singapore. Heterozygous crosses were performed starting at the F2 generation to produce WT, Hets and zygotic KO larvae. These were housed at a density of 50 per tank. 15-20 animals were randomly selected and genotyped every week by scaling to track Mendelian representation over time under conditions of unsegregated growth. Briefly, scales were lysed in 50mM NaOH at 95°C for 15 mins followed by neutralization with 1/10^th^ volume Tris-HCl pH 8.0 and debris pelleted by centrifugation at 3000 g for 5 mins before PCR with LHW4_zfBr_5’_F and LHW7_zfBr_3’_R primers listed above. To enable segregated growth, larvae were genotyped at 3dpf by clipping the tip of the tail fin ^67^ and housed at a lower density of 20/tank. KO fish were given one extra feed per day to boost their growth. The standard length of these larvae were measured once every week under a stereoscope (up to day 28) or with a ruler after minimal tricaine anesthetization. *Br* KO fish that survive past 4 months have a short mating span of about 2 months to generate maternal zygotic (mz KO) larvae. Beyond that point, we notice that the *Br* KO males fail to display effective mating behavior due to their low body weight and lethargic phenotype. Female *Br* KO females are however, still able to mate with WT or Het fish, demonstrating preservation of their fertility.

### Birefringence Analysis

4-6 dpf larvae were anaesthetized briefly with tricaine before embedding in 30% methylcellulose in the egg water. Imaging was performed by placing the dish containing the larvae in between a pair of polarized filters (50mm Diameter Unmounted, Linear Glass Polarizing Filter, Stock No. #43-787 from Edmund Opticals) aligned at 90 degrees to each other. Images were collected under brightfield with a Nikon SMZ800 stereoscope connected to a CCD camera.

### Zebrafish Motility Monitoring

5 dpf mz KO and WT larvae were gently pipetted into a 48-well plate, each fish per well for monitoring locomotor activity. The 48-well plate was placed in a sound-attenuated incubator to isolate it from extraneous noise and light. Inside the incubator, a while-light LED box was positioned below the plate and videos were recorded from the top at a speed of 5.3 frames per second (5.3 fps) by a Basler USB3 camera in a resolution of 2048 × 1024 pixels. The position (x-y coordinates) of the fish was determined in real-time based on background subtraction algorithms written in Python utilizing OpenCV library. This data was then exported into Microsoft Excel files and analyzed offline for speed and distance measures.

### Zebrafish Respiratory Analysis

Manually dechorionated 24 hpf (1 dpf) mz KO larvae were placed into each well (1 per well) of an Agilent XF96e Spheroid microplate containing 180 ul of seasalt water and allowed to adapt for 1 hour. Basal respiration was recorded on the Seahorse XF96e at 28°C. This was achieved by placing the XF96e in a refrigerated chamber held at 16°C. We were unfortunately unable to perform a Mito Stress test on live larvae because the repeated mixing steps following injection of Oligomycin and FCCP caused significant lethality to the larvae due to the low height clearance of the XF96e oxygen probe.

### Zebrafish Metabolomic Analysis

At 5 dpf, 50 WT or mz KO larvae were collected into a tube under normal conditions, placed on ice for 5 mins to immobilize larvae, followed by gentle sedimentation. All egg water was removed and larvae were rinsed washed twice with ice cold water. The dried pellet was weighed to 2 decimal points in mg and snap frozen, thawed on ice and homogenized in 50% H_2_O/50% Acetonitrile using zirconia beads in a liquid nitrogen cooled rotor (2 cycles of 6500 rpm for 20 s). For organic acid extraction, 300 μL of tissue homogenate was extracted with ethylacetate, dried and derivatized with N,O-Bis(trimethylsilyl)trifluoroacetamide, with protection of the alpha keto groups using ethoxyamine (Sigma Aldrich, USA). The mixture was allowed to equilibrate for 30 s and 1.2 ml of HPLC grade methanol was added to the mixture. After vortexing, the mixture was incubated in 50 °C for 10 min and centrifuged to pellet the precipitated protein. The supernatant was removed and dried under nitrogen gas in a clean microcentrifuge tube. The dried extract was reconstituted in 100 μl of methanol before analyzing using a liquid chromatography-mass spectrometer. Trimethylsilyl derivatives of organic acids were separated by gas chromatography on an Agilent Technologies HP 7890A and quantified by selected ion monitoring on a 5975C mass spectrometer using stable isotope dilution. The initial GC oven temperature was set at 70 °C, and ramped to 300 °C at a rate of 40 °C/min, and held for 2 min.

### MitoCore Modeling

The Mitocore model was downloaded in SBML format from^27^. Each of the 485 reactions in the model was blocked (setting upper and lower bounds of permissible flux to zero), one reaction at a time, and the ATP production (Reaction ID: “OF_ATP_MitoCore”) was constrained to have a flux > 90% of the theoretical capacity of the blocked model. Uniform sampling was performed for each blocked model using the CHRR algorithm of Cobra toolbox 3.0 ^68^ to obtain 5000 flux vectors that represent the solution space. As baseline comparison, the unchanged MitoCore model (*unconstrained model)* was also sampled to obtain 5000 flux vectors. The flux distributions for the export reactions of Lactate (Reaction ID:‘L_LACt2r’) and Succinate (Reaction ID: ‘SUMt_MitoCore’) were compared between the *unconstrained model* and each of the 485 constrained models using the Wilcoxon ranksum test, and the effect sizes were computed using Cohen’s d. Reaction blockages that produced a negative effect size for either lactate or succinate export were discarded, and the remaining blocked models were ranked by the sum of effect sizes (for Lactate and Succinate). A similar workflow was repeated to identify which of the 371 genes in the MitoCore model had largest effect size to increase succinate and lactate export when knocked out (Single Gene Deletion studies using Cobra Toolbox 3.0).

### Mitochondria Flux assays

Respirometric analysis of purified mitochondria from zebrafish skeletal muscle was performed using Agilent’s Seahorse platform as described by Boutagy et al. ^69^. Briefly, 1.0-1.4 μg of purified mitochondria were plated in each well of a XF96e cell culture plate. To measure uncoupled electron flow, assay buffer contained 5 mM pyruvate, 4 μM carbonyl cyanide 4-(trifluoromethoxy) phenylhydrazone (FCCP), 1 mM malate. Rotenone (2 μM, final), succinate (5 mM, final), antimycin A (4 μM, final), TMPD/ascorbic acid (100 μM/10 mM) were injected sequentially during the assay. In coupling assay with pyruvate and malate, assay buffer contained 10 mM pyruvate and 5 mM malate. In coupling assay with Complex I inhibited, assay buffer contained 10 mM succinate and 2 μM rotenone. In coupling assay, ADP, oligomycin A, FCCP and antimycin A were injected into assay plate sequentially to 3.2 mM, 3 μM, 4 μM and 4 μM, respectively. Citrate synthase activity (described below) was used for post-normalization of OCR rates.

### Mitochondria Purification from Cultured Cells and Tissues

Mitochondria were isolated from cultured cells using mitochondria isolation kit (Abcam, ab110168). Cells were harvested and ruptured with a Dounce homogenizer in the buffer A containing cOmplete protein inhibitor cocktail (Roche) and 1mM phenylmethylsulfonyl fluoride (PMSF). 15 stokes were performed with a motorised pestle operated at 300 rpm. Nuclei and unbroken cells were removed by centrifugation at 600 g. Homogenates were spun at 7000 g for 15 min to precipitate mitochondria. Mitochondria pellets were collected for downstream uses. To isolate mitochondria from zebrafish, skeletal muscles of fish were dissected and minced. Tissue was disrupted in muscle mitochondria isolation buffer (67 mM sucrose, 50 mM KCl, 1mM EDTA, 0.2% fatty acid free BSA, 50 mM Tri-HCl, pH7.4) with a Dounce homogenizer tight pestle operated at 1300 rpm. The extract was centrifuged at 600 g for 5 min to clear intact myofibrils and heavy cell debris and then spun at 7000 g for 15 min to isolate the desired mitochondrial fraction.

### Immunofluorescence

Cells were fixed by 4% PFA at 37 °C for 20 min and washed with PBS twice. Where appropriate, cells were stained with 25 nM of Mitotracker Red at 37 °C for 30 min before the fixation. Samples were solubilized with 0.3% Triton X at room temperature for 10 min and washed twice with PBS. Cells were blocked with blocking buffer (3% BSA, 10% FBS in PBS) at room temperature for 1 h. Primary antibodies were diluted to desired concentrations in the blocking buffer and incubated with cells for over-night, rocking at 4 °C. Proteins of interest were probed with the following antibodies: anti-BR, 1:200 (Novus, NBP1-90536); anti-HA, 1:500 (BioLegend, 901502); anti-CYTC, 1:500 (Santa Cruz, sc-13561); anti-CS, 1:500 Santa Cruz, SC-390693); anti-TOMM20, 1:500 (Proteintech, 11802-1-AP). Cells were washed with 3 changes of PBS-Tween 20 0.1% (v/v) before incubating with 4 μg/ml of secondary antibodies for 1 h at room temperature (Alexa fluro 488, 594, 647, Invitrogen). Samples were imaged on an Olympus FV3000 confocal laser scanning microscope.

### Antibody Generation

Custom polyclonal antibodies were generated against human BR (hBR) and zebrafish BR (zBr) using the following peptide antigens.

For SDS-PAGE, BN-PAGE, IF and IHC of human/mouse BR: 5’EVVHRYYRPDLTIPEIPPKRGELKTELLGLKERKHKPQVSQQEELKC3’

For SDS-PAGE of zBr: 5’RPDLSIPEIPPKPGELRTELLGLKERQMDSQKQ3’ For BN-PAGE of zfBr: 5’RPDLSIPEIPPKPGELRTEC3’

Peptides were conjugated to KLH and used for rabbit immunization according to standard procedures.

### Blue-Native PAGE analysis

BN-PAGE analysis was performed as described by Jha et al. ^70^ and 2^nd^ dimension BN-SDS PAGE as described by Fiala et al. ^71^ with minimal modifications. All mitochondria were solubilized with 8 g/g of digitonin.

### Immunoblotting

Whole cell or mitochondrial samples were lysed in RIPA buffer supplemented with cOmplete protein inhibitor cocktail (Roche) and 1mM phenylmethylsulfonyl fluorid (PMSF). When appropriate, protein concentration was measured by bicinchoninic acid (BCA) assay (Thermo Fisher). Lysates were boiled in 1X laemmli sample buffer (50 mM Tris-HCl pH6.8, 2% SDS, 10% glycerol, 12.5 mM EDTA, 0.02% bromophenol blue, 50 mM DTT). After resolving by SDS-PAGE in NuPAGE MES SDS running buffer (Thermo Fisher), proteins were transferred to a 0.2 μM PVDF membrane (Thermo Fisher) and incubated in blocking buffer (3-5% non-fat milk TBS-Tween 20 0.1% (v/v)) for 1 h at room temperature. Membranes were probed with appropriate antibodies in TBST milk for overnight, rolling at 4°C. Following antibodies were used: anti-BR, 1:3000 (Novus, NBP1-90536); custom anti-human BR (1 μg/ml); custom anti-zebrafish BR (1 μg/ml); anti-ACTIN, 1:5000 (Sigma, A2228); anti-TOMM20, 1:5000 (Proteintech, 11802-1-AP); anti-CS, 1:2000 (Santa Cruz, SC-390693); anti-ATP5A, 1:5000 (Santa Cruz, SC-136178); anti-TOMM70, 1:5000 (AbClonal, A4349); anti-VDAC1, 1:2000 (AbClonal, A0810); anti-MTCOI, 1:3000 (Abcam, ab14705); anti-TIM23, 1:5000 (Proteintech, 11123-1-AP); anti-OXPHOS cocktail, 1:5000 (Invitrogen, 45-7999); anti-UQCRC1, 1:5000 (AbClonal, A3339); anti-NDUFA9, 1:3000 (AbClonal, A3196); anti-SDHA, 1:5000 (AbClonal, A2594); anti-TUBULIN, 1:5000 (Sigma, T9822); anti-pACC, 1:2000 (CST, 3661S); anti-GAPDH, 1:5000 (CST, 2118S); anti-FLAG, 1:2000 (BioLegend, 637302); anti-C4ORF48 (Abcam, ab185315); anti-COX4, 1:3000 (AbClonal, A10098). Membranes were extensively washed with four exchanges of fresh TBST and probed with HRP conjugated secondary antibodies against mouse, rabbit or rat IgG (Jackson ImmunoResearch) and washed again. Chemiluminescence was captured by X-ray films (Santa-Cruz) or a Chemidoc imaging system (Bio-Rad).

### Sodium Carbonate Extraction

Sodium carbonate extraction was performed as described with minor modifications ^72^. Mitochondria pellets isolated from HEK293T cells were resuspended in buffer C (320mM Sucrose, 1mM EDTA, 10mM Tris-Cl pH 7.4), buffer C containing 1% Triton X or 0.1M Na_2_CO_3_ pH11. The resuspensions were incubated at 4°C for 1 h with gentle nutation. Supernatant and insoluble pellet were separated by centrifugation at 20, 000 g for 30 min, and the supernatant was gently removed without disturbing the pellet. Equal fractions of supernatant and pellet were analysed by SDS-PAGE immunoblotting.

### Mitochondrial Membrane Fractionation

Mitochondria were isolated with isotonic buffer (250 mM sucrose, 1mM EDTA, 10 mM HEPES-KOH pH7.4, protease inhibitor cocktail and 1 mM PMSF) from HEK293T cells following procedures described in the mitochondria isolation section. Mitochondria pellet was gently resuspended in the isotonic buffer to get protein concentration at 1 mg/ml. Aliquots of mitochondria were solubilized by increasing concentrations of digitonin (0%, 0.1%, 0.115%, 0.13%, 0.145%, 0.16%, 0.175%, 0.19%, 0.205%, 0.22%) for 1 h at 4 °C. Soluble and insoluble fractions were separated by centrifugation at 20, 000 g for 20 min. Insoluble fractions were fully resuspended in an equal amount of isotonic buffer to match the volume of supernatant. Both fractions were lysed in 1X laemmli sample buffer and analysed by SDS-PAGE and immunoblotting.

### Protease Sensitivity Assay

Mitochondria were isolated with isotonic buffer (250 mM sucrose, 1mM EDTA, 10 mM HEPES-KOH pH7.4, protease inhibitor cocktail) from HEK293T cells. Mitochondria pellet was washed to eliminate protease inhibitors. Mitochondria were resuspended to a protein concentration of 1 mg/ml. Mitochondrial membranes were by differentially solubilized by increasing concentrations of digitonin (0-0.2% as indicated in the figure) or 1% Triton X for 10 min on ice. Proteinase K was then added to 100 μg/ml and incubated for 30 min on ice to allow the complete digestion of accessible proteins. To terminate the protease digestion, PMSF was freshly prepared and added to a concentration of 8 mM. Samples were analysed by western.

### BR Interactome Analysis

To determine physical interaction partners of BRAWNIN (BR), we purified BR (non-tagged and FLAG tag) with different antibodies at different expression levels (endogenous and over-expressed). To purify endogenous BR, mitochondria were isolated from HEK293T cells as described. Mitochondria were solubilized in 1 ml of IP buffer (150 mM NaCl, 1 mM EDTA, 50 mM HEPES-KOH pH 7.4) containing 1% n-Dodecyl β-D-maltoside (DDM), cOmplete, EDTA-free protease inhibitor cocktail (Roche) and 1 mM PMSF at 4 °C for 30 min with constant rocking. Homogenate was spun at 20, 000 g for 10 min at 4 °C to remove insoluble content and the supernatant was collected and incubated with BR antibody (Novus, NBP1-90536) or Rabbit IgG control for overnight. 30 μl of protein A/G beads were washed with IP buffer containing 0.1% DDM for two times and incubated with the protein lysate for 2 h at 4 °C to allow antibody/protein complex binding. Non-interacting proteins were removed by four rounds of spin and IP buffer washes in the presence of 0.5% DDM. The beads were then washed with IP buffer only. Bound proteins were eluted with 0.1% Rapigest at room temperature. For the isolation of over-expressed BR, a similar workflow was performed except that BR was transiently over-expressed in HEK293T before the sample preparation. For isolating FLAG tagged BR, we fused a single FLAG tag to the c-terminus of BR. BR and BR-FLAG were transiently overexpressed in HEK293T. Mitochondria were isolated from the two samples separately and solubilized with IP buffer (150 mM NaCl, 1 mM EDTA, 50 mM HEPES-KOH pH 7.4) containing 1% n-Dodecyl β-D-maltoside (DDM), cOmplete, EDTA-free protease inhibitor cocktail (Roche) and 1 mM PMSF. After clearing the insoluble fraction by spin, protein lysates were incubated with anti-FLAG M2 resin (Sigma) for overnight with nutation. The resin was washed as described above and eluted with 0.1% Rapigest (Waters).

To prepare for mass spectrometry, samples were treated with 100 mM triethylammonium bicarbonate (TEAB) pH 8.5 and 20 mM tris(2-carboxyethyl)phosphine hydrochloride TCEP for 20 min at 55 °C with shaking. Proteins were then alkylated by 55 mM chloroacetamide (CAA) at room temperature for 30 min in dark. 3 volumes of 100 mM TEAB were added to dilute out Rapigest to allow following protease digestion. The sample was first treated by 2.5 μg of LysC (Wako) for 3 h at 37 °C with shaking and then treated with 2.5 μg of Trypsin (Promega) for over-night at 37 °C with shaking. Rapigest was hydrolysed by adding TFA to 1% and removed by spin at 20, 000 g for 10 min. The sample was desalted on a C18 cartridge (Oasis) and eluted with 50% ACN and 100 mM TEAB pH8.5, vacuum concentrated and resuspend in 100 mM TEAB pH 8.5. Peptides were labelled with 10 plex tandem mass tags (TMT) for over-night and then quenched by 0.5M Tris-HCl pH7.4, pooled, desalted again and analysed by an Orbitrap LC/MS as described^73^.

### BR Expression by Adeno-associated Virus

BR Adeno-associated Virus (AAV) expression construct was generated by inserting the murine BR ORF into a pscAAV2 backbone with a single FLAG peptide fused to the c-terminus of BR. AAV9-pseudotyped virus was produced by Vector Core, Genome Institute of Singapore, Singapore. 10^9^ viral genomes were constituted in 10 μl PBS and administrated. Tissues were isolated 1 month after the AAV administration.

### Cell Culture

All cell lines were obtained from American Type Cell Collection (ATCC; www.atcc.org) and cultured using standard tissue culture techniques. HEK293T (ATCC® CRL-3216™) cells were cultured in Dulbecco’s High Glucose Modified Eagles Medium/High Glucose (HyClone). U87MG (ATCC® HTB-14™) cells were cultured in Minimal Essential Medium with L-Glutamine (Gibco). All media were supplemented with 10% South American sourced Fetal Bovine Serum (HyClone) and 1% of 10,000U/mL of Penicillin-Streptomycin (Gibco). Starvation regimes were performed as follows: HEK293T cells were plated in normal DMEM (4500mg/L glucose) overnight, followed by a complete media change to that containing 3 μM Etomoxir in complete media, media lacking FBS (Serum-) and or DMEM containing 100mg/L of glucose (GLUCOSE-).

### BR Over-expression by Lentivirus Transduction

The human BR ORF was inserted into a pCDH lentivirus vector under the control of a CMV promoter. Plasmid was then packaged into viral particles using the pPACKH1 lentivector Packaging Kit (System Bioscience) in HEK293T cells. 25% of total crude virus extracted was used to infect U87MG for 2 days before G418 (Santa Cruz) selection at 1.2mg/mL.

### Brawnin Knockdown

siRNA-mediated knockdown of *BR* was performed in U87MG with ON-TARGETplus Human C12orf73 (728568) siRNAs (Dharmacon) with the following sequences : 5’CAGUGGAUGUUUAGCCGAU3’ (siBR_14) and 5’CGACCGGACCUGACAAUAC3 (siBR_16). An equimolar ratio of both targeting siRNAs are mixed to achieve maximal knockdown. Control siRNA sequence is 5’UGGUUUACAUGUCGACUAA3’. Briefly, siRNAs are transfected at 25nM final concentration using Dharmafect using a suspension transfection protocol for U87MG.

shRNA-mediated knockdown or *BR* was performed using the following sequences: 5’AAAAGTTCGCGGTACAGTCTCTAGT3’ (shBR_1) and 5’

AAAAGTGGATGTTTAGCCGATACGT3’ (shBR_3). shRNA hairpins were cloned into a lentiviral backbone to allow stable expression. Viral particles were produced in LentiX293 and purified according to standard procedures. 500,000 U87MG cells were collected and resuspended with respective medium with 32μg/mL of polybrene, and added to polybrene-treated cells at a target MOI of 3. Cell-virus suspension solution was centrifuged at 1000g for 1 hour at room temperature to maximize the collision between cells and viruses. The mixture was then diluted 4 times with respective medium and seeded to one 6-well plate (Costar). Following recovery from viral transduction, cells were selected with 1ug/ml of puromycin until a non-transduced control showed complete death. U87MG Scr, shBR cells were used within 10 passages of initial transduction to minimize phenotypic drift due to compensation.

### Seahorse Cultured Cell MitoStress Test

15,000 U87MG cells were plated on poly-L-Lysine (Sigma) treated Seahorse XF96 Cell Culture Microplates (Agilent) and XFe96 sensor cartridge (Agilent) was hydrated with distilled water at 37°C with no CO_2_ the day prior to MitoStress Assay. On the day of actual MitoStress Assay, cultured cells were washed once and media were replaced with XF basal DMEM supplemented with 1 mM pyruvate (Sigma), 2 mM glutamine (Sigma), and 10 mM glucose (Sigma). Pre-hydrated sensor cartridge was calibrated with Seahorse XF Calibrant (Agilent). Both cell culture plate and sensor cartridge were incubated at 37°C with no CO_2_ for at least 45 mins before actual assay starts. For MitoStress assay, 2 μM of oligomycin (Sigma), 1 μM of FCCP (Sigma), and 1μM of Rotenone/Antimycin (Sigma/Sigma) were injected according to Seahorse MitoStress Assay Protocol (Agilent). Citrate Synthase Normalization Assay was performed on the cultured cell plate to normalize the measured Oxygen Consumption Rate (OCR) and ExtraCellular Acidification Rate (ECAR).

### Citrate Synthase Normalization Assay

Citrate Synthase Normalization Assay was performed on Seahorse XF96 Cell Culture Microplate (Agilent) right after seahorse assay or on plates frozen at −80°C with leftover medium aspirated. All leftover medium was removed and replaced with 113 μl of CS buffer (200mM Tris buffer at pH8.0, 0.2% Triton X-100 (v/v), 100μM DTNB (Sigma) in 100mM Tris buffer at pH8.0, 1mM Acetyl-CoA (Sigma)) per well. 5μl of 10mM Oxaloacetate (Sigma) in water was added to each well as reaction substrate. Absorbance at 412nM at 37°C was recorded with Tecan Microplate Reader M200 (Tecan) at minimal time interval for 8mins. Citrate Synthase activity was then calculated using the formula below

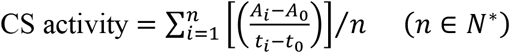

n is the total number of absorbance records; A is the absorbance recorded; t is the time.

### Respiratory Chain Enzymatic Assays

Respiratory chain enzymatic assays were performed with zebrafish skeletal muscle homogenates according to the protocol as described by ^28^.

### Statistical Analysis

Unless otherwise stated, all data are presented as Mean ± standard error of mean (SEM.). Statistical approaches to test differences of means were performed using Prism 5.0. The statistical test used in each panel is indicated in each accompanying figure legend. Where p values are not explicitly indicated, significance levels follow the convention of *p<0.05, **p<0.01, ***p<0.001.

## Acknowledgements

We thank the IMCB Aquatics facility, the IMU Imaging facility and the AMPL histopathology unit at A*STAR for providing technical support; **JP Kovalik** and **Ching Jianhong** from the Duke-NUS metabolomics facility for metabolomic analysis including assay development; **Hu Zhen, Kor Chia Yee** and **Jes Kwek Hui Min** for their technical support; **Aida Moreno Moral** and **Cheryl Lee** for their inputs on bioinformatics analyses; **Brijesh Kumar Singh** for his assistance with the Agilent Seahorse platform; **Sudipto Roy** from IMCB for help with zebrafish muscle analyses. Lastly, we are extremely grateful to **Wong Pui Mun** for performing initial experiments to test the hypothesis that BR is a secreted peptide, **Serene Chng** for guidance on zebrafish crispr knockout generation.

## Author contributions

L.H. and S.Z. conceptualized the study, designed, performed and interpreted all experiments. S.Z., G.F. carried out the SEP screen. L.C, P.J.H., C.T., G.F., R.C., S.J., J.F. and C.W. performed and interpreted specific experiments. B.K., V.O., G.M. and E.P. designed and performed computational analyses on sORF selection and mitochondrial functional prediction. S.N., P.S., B.R. and S.L. provided reagents. R.S., L.C.W., and D.S. carried out proteomic analyses. C.M. carried out experiments and P.D.R. performed conservation analyses under the supervision of L.L. N.S.J and L.T.K. implemented MitoCore model. D.S. performed proteomics and biochemical analyses and edited the manuscript. L.H. and S.Z wrote the manuscript.

## Data and materials availability

All data will be made available upon publication. All R code for WCGNA/GSEA analyses available upon request. Proteomics data will be deposited in Proteome Xchange upon publication. All reagents cited, including animals, will be available upon request through MTA.

