## Supplemental Information for "BRAWNIN: A sORF-encoded Peptide Essential for Vertebrate Mitochondrial Complex III Assembly"

**This PDF file includes:**

Supplementary Text
Figs. S1 to S4
Table S5
Supplemental References

**Other Supplementary Information for this manuscript include the following:**

Excel files of Table 1-4

**Supplementary Text**

*sORF selection Parameters*

To determine the quality and likelihood of each sORF, multiple algorithms have been developed to assess ribosomal periodicity and control for inherent biases in the Ribo-Seq workflow.

1) ORFscore [1]: The ORFscore calculates the preference of ribosome protected fragments (RPFs) to accumulate in the first frame of coding sequences. To compute the ORFscore, RPFs are counted in each frame (in - frame 0, +1 frame and +2 frame). Next, the distribution of RPFs is compared to an equally sized uniform distribution using a modified chi-squared statistic. In literature, an ORFscore of at least 6 is generally considered a good score.

2) In-frame-coverage (IFC) [2]: Percentage of nucleotides covered by in-frame situated RPFs (i.e. mapping at first base of coding triplets to the last base of the stop codon).

3) Coverage Uniformity (U) [2]: Represents how uniform the ribosome footprints are distributed over the sORF sequence. This filter ranges between -1 and 1, with either boundary indicating that all ribosomes reside in one half of the sORF. A coverage uniformity of 0 implies that the ribosomes are uniformly distributed.

4) FLOSS [3]: The FLOSS algorithm is an additional method designed to distinguish true coding from non-coding sequences based on the RPF length distribution. The FLOSS algorithm provides a score based on the comparison between the RPF length distribution in each sORF and the RPF length distribution found in canonical protein-coding sequences. Based on the FLOSS score, a classification is made representing the coding tendency of sORFs.

*Mitochondrial Prediction*

The first method relies on mitochondrial functional prediction based on functional enrichment of SEP co-expressed genes. First we performed co-expression analysis where SEPs were ranked by weighted gene correlation network analysis (WGCNA) of their host gene with genes in publicly available human liver, heart and skeletal muscle RNAseq datasets (Fig. S1a, details in Method section). Next, we performed functional enrichment analysis using gene set enrichment analysis (GSEA) of ranked lists from WGCNA. To classify each candidate as mitochondrial or non-mitochondrial, we performed multi-dimensional PCA analysis of pathway normalized enrichment scores (NES) of MitoCarta ("mito") genes and a matched number of randomly chosen "non-mitochondrial" transcription factors. We used non-hierarchical K-means clustering, averaging over 15 clusters to obtain a mitochondrial likelihood score of each SEP candidate in each of the 3 tissues (Fig. S1a). The threshold defining mitochondrial identity in each tissue was set by maximising the predictive performance of 53 known MitoCarta SEPs against 83 known non-mitochondrial SEPs. Using this method, 64 "high confidence" SEPs were included because they scored positive in all of the tissues in which they are expressed and have not been previously characterized (Fig. S1a). This method correctly classified 73% of known training set SEPs with a false positive and negative rate of 6% and 57% respectively. Notably, the classification success drops to 60% when performed in cultured skin or endothelial cells, which have lower mitochondrial content compared to heart, muscle and liver (data not shown).

Next, we selected 72 SEPs predicted to be mitochondrial because they contained a potential "mitochondrial targeting motif". This motif can be either a transmembrane domain (TM) or signal peptide (SP) with reduced hydrophobicity, a mitochondrial targeting sequence (MTS) or twin-cysteine (2C3XC, 2C9XC, C9XC & C10XC). This combination correctly retrieves 87% of known MitoCarta SEPs but has a very high false positive rate of 49% (based on known non-mitochondrial SEPs).

The third method relies on empirical evidence of protein expression in the mitochondria. 23 thresholded SEPs with matching spectra in an LC-MS/MS dataset of purified human mitochondria [4] (details in the Method section) with unknown function were included regardless of their score in the first two measures.

*SEP Screening Outcome*

Of 173 SEP candidates tested, we were able to obtain a credible HA fluorescence signal from the overexpression of the open reading frames of 88 candidates (50.9%) in HeLa (Fig. S1c), representing the experimental validation of 88 novel sORF-encoded peptides (SEPs). SEP candidates with undetectable expression levels may be unstable, require the presence of autologous UTRs for transcript stability or translational efficacy, or might be secreted upon production. These were excluded from further characterization. Notably, the highest rate of successful protein expression was from the category of annotated sORFs (88.2%). 48.5% uORF and 37.5% of lincRNA-derived SEPs can also generate stable protein. Of the 3 approaches used for mitochondrial prediction, the strongest predictor was the presence of protein motifs, namely a transmembrane domain (TMD), with 30.5% positive prediction rate (Fig. S1d). WGCNA/GSEA had a poorer-than-expected outcome because 82.8% of GSEA-predicted candidates were uORF-derived SEPs. Since GSEA utilizes the underlying host gene for functional prediction, which cannot distinguish between the main annotated ORF and the uORF-encoded proteins, we are unable to accurately decipher the function of the uORF-derived SEP. Lastly, the presence of matching spectra in existing proteomics dataset turned out to be the poorest predictor for protein validity, potentially due to off-target identifications. This underscores the importance of unbiased, experimentally-validated screening efforts as reported in this study. By inference, the presence of matching spectra alone should not be used as definitive evidence of the validity of a SEP. Of note, validated SEPs that do not localize to the mitochondria display a wide variety of subcellular localization (Fig. S1e), suggesting their potentially diverse functions in cell biology.

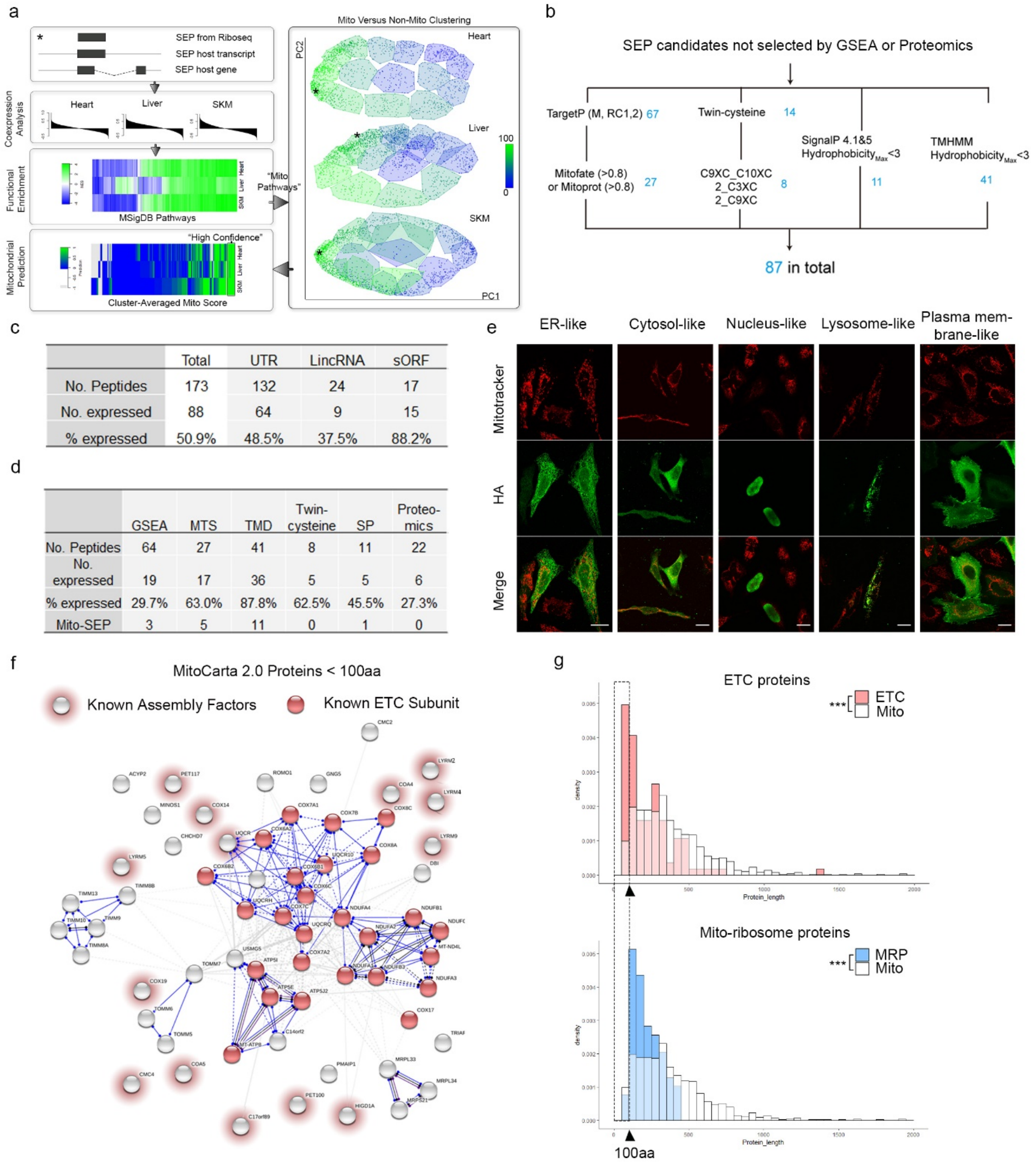


| | Total | UTR | LincRNA | sORF |
|---|---|---|---|---|
| No. Peptides | 173 | 132 | 24 | 17 |
| No. expressed | 88 | 64 | 9 | 15 |
| % expressed | 50.9% | 48.5% | 37.5% | 88.2% |

| | GSEA | MTS | TMD | Twin-cysteine | SP | Proteo-mics |
|---|---|---|---|---|---|---|
| No. Peptides | 64 | 27 | 41 | 8 | 11 | 22 |
| No. expressed | 19 | 17 | 36 | 5 | 5 | 6 |
| % expressed | 29.7% | 63.0% | 87.8% | 62.5% | 45.5% | 27.3% |
| Mito-SEP | 3 | 5 | 11 | 0 | 1 | 0 |

**Figure S1. Mito-SEP prediction pipeline identifies novel endogenous mitochondrial SEPs**

a. Co-expression analysis coupled to functional enrichment analysis to identify "high confidence" SEP genes with mitochondrial gene signature. Color bar scale on the right panel refers to the percentage of known mitochondrial genes in each K-means cluster. Color bar scale on the bottom left panel refers to the K-means score. All other color bars depict normalized enrichment scores

(NES). *indicates a specific “high confidence” candidate as it transits through the pipeline. See SI text for details.

b. Workflow for identifying Mito-SEPs with potential mitochondrial targeting protein domains.

c. Numbers of screened and successfully expressed peptides (i.e. with detectable HA IF signal) according to their biotype annotation as 5’UTR, LincRNA or sORF in their genomic loci.

d. Numbers of screened and expressed peptides that were predicted by indicated methods. Gene Set Enrichment Analysis (GSEA), Mitochondrial Targeting Sequence (MTS), Trans-membrane domain (TMD), Signal Peptide (SP).

e. Representative SEP candidates with validated protein expression by HA IF, but that are not mitochondrial. Different categories of subcellular localization observed are shown and described here. Scale bar = 20 µm.

f. MitoCarta 2.0 proteins <100 a.a. were analyzed by STRING V10 [5] and classified according to functional ontology (GO and manual curation). Dark red circles refer to proteins with known functions in electron transport chain (ETC) complexes; light red halos refer to proteins with known role as ETC assembly factors.

g. Upper panel: Size distributions of ETC proteins (complex subunits and assembly factors) and all mitochondrial proteins (Mitocarta 2.0). Lower panel: Size distribution of mitochondrial ribosome proteins (MRP) and all mitochondrial proteins (Mitocarta 2.0). The statistical difference of their size distributions were tested by Mann-Whitney U-test. Note the relative enrichment of ETC proteins that are below 100 a.a., compared to mito-ribosome proteins, even though components of both complexes are enriched for lower molecular weight proteins.

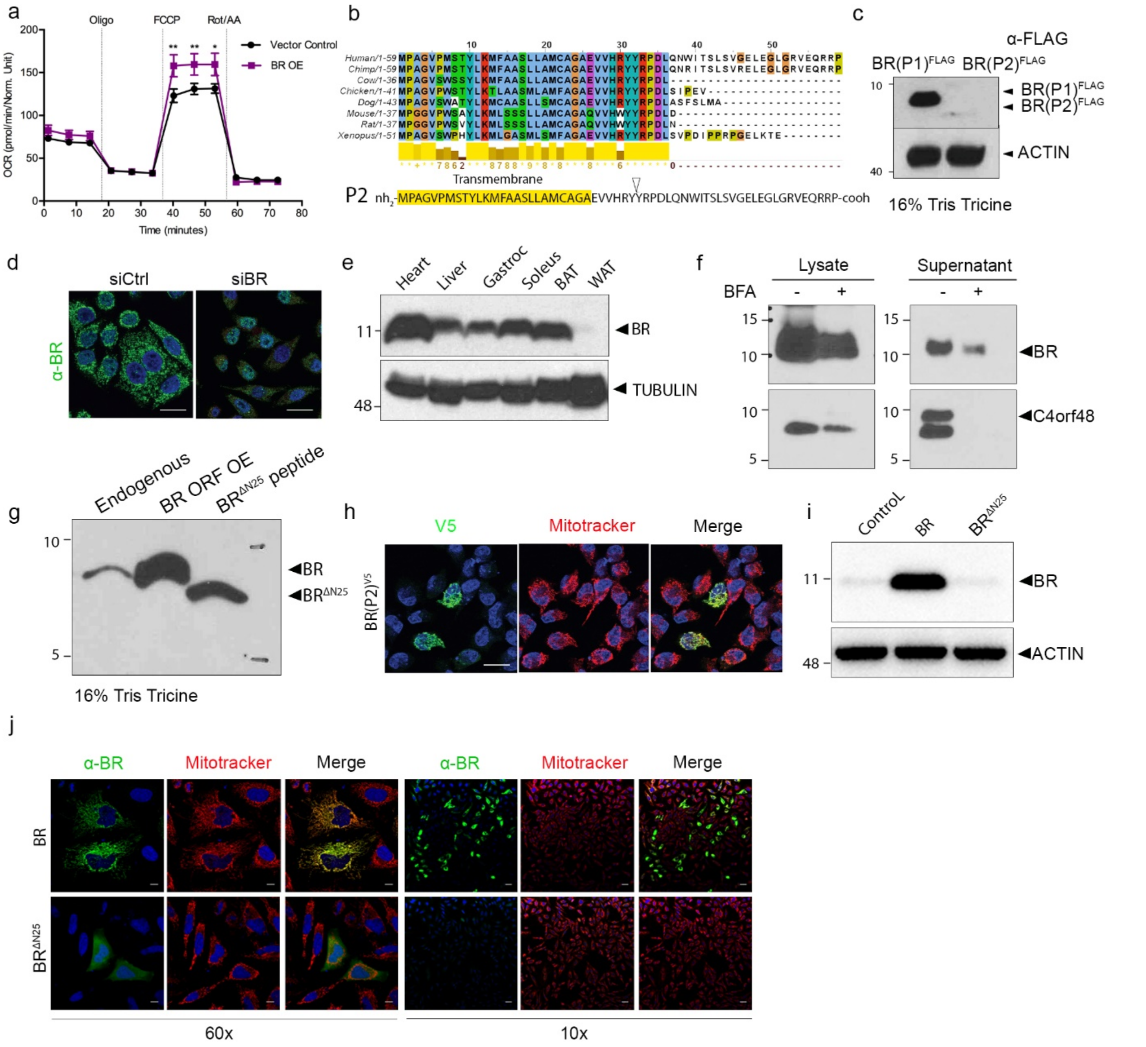


**Figure S2. BRAWNIN (BR) is a conserved SEP at the inner mitochondrial membrane**

a. Seahorse MitoStress test of U87MG cells stably transduced with a lentiviral construct expressing BR or empty vector. Oligo, oligomycin; FCCP, carbonyl cyanide 4-(trifluoromethoxy)phenylhydrazone; Rot/AA, rotenone/antimycin A.

b. Alignment of the predicted BR isoform 2 (BR-P2) encoded by an alternatively spliced transcript of the *C12orf73* gene.

c. Western blot of lysates from HEK293T transfected with BR isoform 1-FLAG (BR(P1)$^{FLAG}$) and isoform 2-FLAG BR (BR(P2)$^{FLAG}$). BR-P2 (59 a.a.) is detectable, but is present at much lower levels compared to BR-P1.

d. IF of endogenous BR in HeLa transfected with control non-targeting and BR-targeting siRNAs.

Scale bar = 20 µm.

e. Western blot of endogenous mouse BR in the indicated tissue lysates. BAT = brown adipose tissue, WAT = white adipose tissue.

f. Western blot of intracellular (lysate) versus secreted (supernatant) BR of HEK293T cells transfected with the BR ORF. C4ORF48, a bona fide secreted peptide of the same length (unpublished) is used as a control. Brefeldin A (BFA) inhibits the classical secretory pathway. Note that BR secretion is not completely abrogated by BFA as would be expected of a bona fide secreted protein.

g. Western blot of endogenous, overexpressed BR in HEK293T separated on a 16% Tris Tricine gel together with a synthetic BR peptide lacking N-terminal 25 amino acids ($BR^{\Delta N25}$).

h. BR-P2-V5 tagged protein can be detected by IF when overexpressed in HeLa and localizes to the mitochondria as indicated by colocalization with MitoTracker. Scale bar = 20 µm.

i. Western blot of HEK293T lysates transfected with full length BR and BR lacking N-terminal 25 amino acids encoding the predicted signal peptide or transmembrane domain ($BR^{\Delta N25}$).

j. IF of transiently over-expressed BR and $BR^{\Delta N25}$ with α-BR in HeLa. Scale bar = 10 µm (60X); 50 µm (10X).

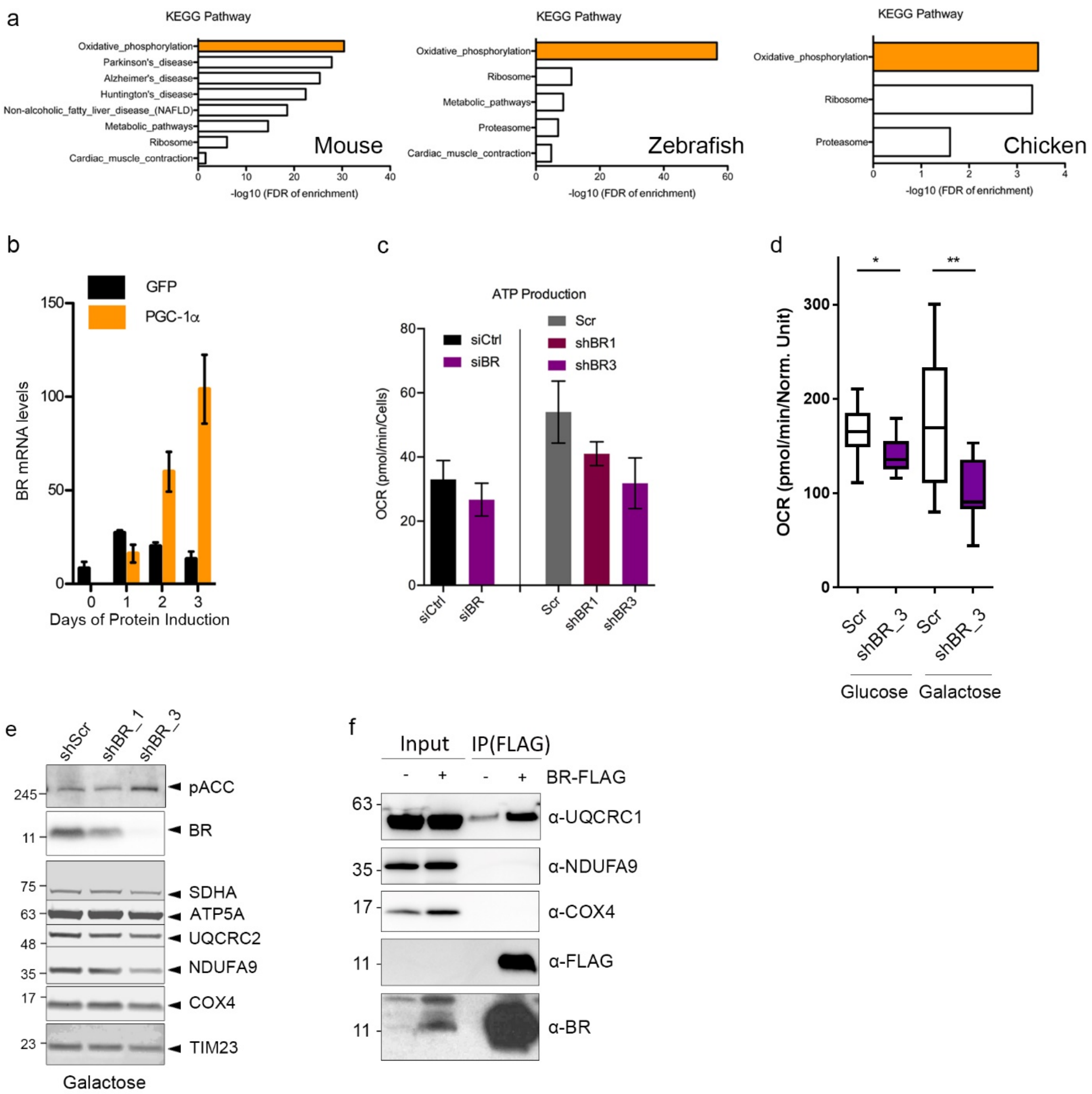


**Figure S3. BR is an AMPK target that potentiates OXPHOS and respiratory complex assembly**

a. Enriched KEGG pathways by Panther gene ontology analysis of the top 200 BR co-expressed genes (from COXPRESdb v7.0 [6]) in mouse, zebrafish and chicken.

b. *Br* mRNA levels in mouse myotubes with enforced expression of PGC-1α. Data are obtained from GEO omnibus dataset GDS1879 with probeset 114297_f_at.

c. ATP production calculated from experiment in Fig. 3d,e. Data are mean and SEM of 2 and 3 biological replicates for shRNA and siRNA-mediated depletion, respectively, each with 6 technical replicates. p-values from unpaired t-test ($\alpha$=0.05).

d. Basal OCR of HEK293T with stable shRNA-mediated knockdown of BR grown under normal (glucose) and oxidative (galactose) conditions. Data are mean and SEM of 6 technical replicates. p-values from unpaired t-test ($\alpha$=0.05).

e. SDS-PAGE and western blot analysis of HEK293T with stable shRNA-mediated knockdown of BR cultured in galactose.

f. FLAG IP followed by Western blotting of mitochondrial solubilized with 1% digitonin from mouse heart injected with Br-FLAG-expressing adeno-associated virus (AAV).

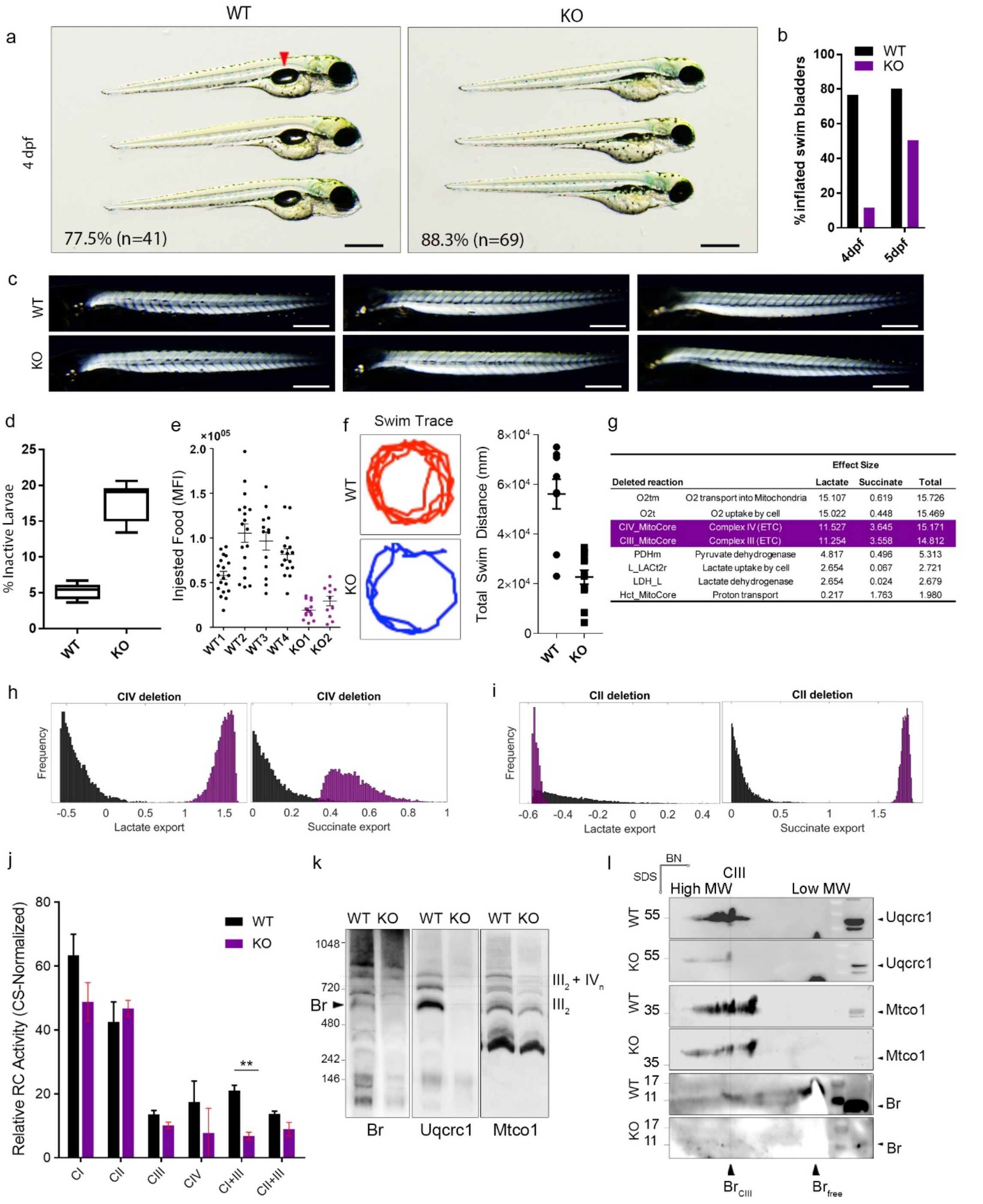


| Deleted reaction | | Effect Size Lactate | Succinate | Total |
|---|---|---|---|---|
| O2tm | O2 transport into Mitochondria | 15.107 | 0.619 | 15.726 |
| O2t | O2 uptake by cell | 15.022 | 0.448 | 15.469 |
| CIV_MitoCore | Complex IV (ETC) | 11.527 | 3.645 | 15.171 |
| CIII_MitoCore | Complex III (ETC) | 11.254 | 3.558 | 14.812 |
| PDHm | Pyruvate dehydrogenase | 4.817 | 0.496 | 5.313 |
| L_LACt2r | Lactate uptake by cell | 2.654 | 0.067 | 2.721 |
| LDH_L | Lactate dehydrogenase | 2.654 | 0.024 | 2.679 |
| Hct_MitoCore | Proton transport | 0.217 | 1.763 | 1.980 |

**Figure S4. Knockout of *Br* in zebrafish causes lethal mitochondrial deficiency**

a. Brightfield images of 4 days post fertilization (dpf) WT and KO larvae. Images represent indicated percentage of all larvae analyzed with indicated n numbers. Swim bladder in WT is indicated by red arrowhead. Scale bar = 100 µm.

b. Quantification of the percentage of larvae from an independent clutch with inflated swim bladders at the indicated age.

c. Birefringence analysis of 5 dpf WT and KO larvae to reveal skeletal muscle fiber integrity and organization. Scale bar = 150 µm.

d. Percentage of inactive mz KO larvae at 6 dpf, scored as larvae that remain motionless on the bottom of the tank for 1 minute.

e. Feeding behavior as measured by the amount of fluorescent paramecium ingested at 7 dpf in 4 clutches of WT and 2 clutches of mz KO larvae. MFI = mean fluorescence intensity of gut region post-feeding. Each dot represents one animal.

f. Representative swim trace of WT versus mzKO larvae at 11 dpf (left). Quantitation of overall distance swam in a fixed assay interval (right).

g. MitoCore reactions, ranked by how strongly their shutdown would increase both lactate and succinate export. (see SI for details).

h. Distributions of lactate and succinate export flux levels in 5000 random samples under unconstrained or CIV-flux-constrained.

i. Distributions of lactate and succinate export flux levels in 5000 random samples under unconstrained or CII-flux-constrained.

j. Respiratory chain enzymatic activities using adult WT and KO skeletal muscle homogenates. Data are mean and SEM of 4 biological replicates. p-values are from paired t-test ($\alpha$=0.05). The difference in CIII activity is just under significance ($p = 0.072$).

k. BN-PAGE and western blot analysis with zebrafish-specific $\alpha$-Br that recognizes native Br. Position of Br is indicated by black arrowhead as this signal is completely absent in the KO. Membrane was completely stripped and reprobed with Uqcrc1 (rabbit polyclonal) and Mtco1 (mouse monoclonal) to reveal the position of CIII and CIV respectively. These data are representative of 4 independent BN-PAGE experiments and further confirmed by 2$^{nd}$ dimension SDS-PAGE described in l.

l. 2$^{nd}$ dimension SDS-PAGE analysis to confirm that the CIII-comigrating signal in (I) is indeed specific to Br, as judged by a signal at the expected 10 kD in the WT which is absent in the KO. Position of CIII is inferred from Uqcrc1 western blotting. Zebrafish Br co-migrates with CIII ($Br_{CIII}$).

**Table S1. Candidates screening information**

sORF screened information : Ensembl gene and transcript ID, origin of prediction, nucleotide and aa sequences and plasmid constructs.

Attached EXCEL sheet Table S1

**Table S2. Gene sets shortlisted for mitochondrial signature per dataset**

List of the gene sets which have a mitochondrial signature (100 mito genes with an absolute NES score above 3) in at least one of the datasets. The columns named after the datasets contain an X when the corresponding gene set was used for the principal component analysis (PCA).

Attached EXCEL sheet Table S2

**Table S3. Protein IDs of BR-interacting proteins detected by IP/MS from 4 independent IP/MS experiments**

The first tab lists protein IDs used for generating venn diagram in Fig. 3g. Information on protein identifications from each experiment is listed in separate tabs.

Attached EXCEL sheet Table S3

**Table S4. Expression correlation between BR and CIII assembly factors**

Ranked of ETC complexes subunits and assembly factors from BR pairwise correlation in different datasets. Ranked percentage is the mean ranked corrected for the genome size of the dataset. (cor = correlation)

Attached EXCEL sheet Table S4

**Table S5. MitoCore: Individual deletions of CIII and CIV genes explain metabolomic observations.**

Top genes ranked by their sum of effect size on lactate and succinate fluxes upon deletion. Highlighted in green are genes from complexes III and IV, in orange are genes from complex I.

| Rank | Gene Symbol | Gene Name | Effect Size Lactate | Effect Size Succinate | Sum |
|---|---|---|---|---|---|
| 1 | UQCRFS1 | ubiquinol-cytochrome c reductase, Rieske iron-sulfur polypeptide 1 | 11.772 | 3.712 | 15.485 |
| 2 | COX4I2 | cytochrome c oxidase subunit 4I2 | 11.759 | 3.653 | 15.412 |
| 3 | CYC1 | cytochrome c1 | 11.718 | 3.689 | 15.408 |
| 4 | COX4I1 | cytochrome c oxidase subunit 4I1 | 11.657 | 3.717 | 15.374 |
| 5 | MT-CO1 | mitochondrially encoded cytochrome c oxidase I | 11.675 | 3.695 | 15.370 |
| 6 | COX8C | cytochrome c oxidase subunit 8C | 11.703 | 3.646 | 15.349 |
| 7 | COX5A | cytochrome c oxidase subunit 5A | 11.679 | 3.635 | 15.314 |
| 8 | CYCS | cytochrome c, somatic | 11.660 | 3.651 | 15.311 |
| 9 | COX6A2 | cytochrome c oxidase subunit 6A2 | 11.699 | 3.606 | 15.304 |
| 10 | COX6C | cytochrome c oxidase subunit 6C | 11.586 | 3.678 | 15.264 |
| 11 | UQCRH | ubiquinol-cytochrome c reductase hinge protein | 11.539 | 3.719 | 15.258 |
| 12 | COX5B | cytochrome c oxidase subunit 5B | 11.541 | 3.683 | 15.223 |
| 13 | COX7A2 | cytochrome c oxidase subunit 7A2 | 11.559 | 3.629 | 15.188 |
| 14 | COX8A | cytochrome c oxidase subunit 8A | 11.495 | 3.656 | 15.151 |
| 15 | TTC19 | tetratricopeptide repeat domain 19 | 11.510 | 3.623 | 15.133 |
| 16 | COX7B | cytochrome c oxidase subunit 7B | 11.480 | 3.633 | 15.113 |
| 17 | UQCRC2 | ubiquinol-cytochrome c reductase core protein 2 | 11.413 | 3.657 | 15.070 |
| 18 | UQCR10 | ubiquinol-cytochrome c reductase, complex III subunit X | 11.454 | 3.614 | 15.068 |
| 19 | COX7C | cytochrome c oxidase subunit 7C | 11.386 | 3.651 | 15.038 |
| 20 | UQCRB | ubiquinol-cytochrome c reductase binding protein | 11.377 | 3.592 | 14.969 |
| 21 | MT-CO3 | mitochondrially encoded cytochrome c oxidase III | 11.311 | 3.641 | 14.952 |
| 22 | COX7A1 | cytochrome c oxidase subunit 7A1 | 11.339 | 3.594 | 14.933 |
| 23 | MT-CO2 | mitochondrially encoded cytochrome c oxidase II | 11.384 | 3.541 | 14.925 |
| 24 | COX7A2L | cytochrome c oxidase subunit 7A2 like | 11.257 | 3.664 | 14.921 |
| 25 | COX6B2 | cytochrome c oxidase subunit 6B2 | 11.203 | 3.597 | 14.801 |
| 26 | UQCR11 | ubiquinol-cytochrome c reductase, complex III subunit XI | 11.308 | 3.492 | 14.800 |
| 27 | COX6A1 | cytochrome c oxidase subunit 6A1 | 11.232 | 3.520 | 14.752 |
| 28 | MT-CYB | mitochondrially encoded cytochrome b | 11.225 | 3.498 | 14.722 |
| 29 | COX6B1 | cytochrome c oxidase subunit 6B1 | 11.018 | 3.567 | 14.585 |
| 30 | UQCRC1 | ubiquinol-cytochrome c reductase core protein 1 | 10.970 | 3.422 | 14.392 |
| 31 | DLD | dihydrolipoamide dehydrogenase | 5.113 | 0.249 | 5.363 |
| 32 | DLAT | dihydrolipoamide S-acetyltransferase | 4.867 | 0.452 | 5.320 |
| 33 | PDHB | pyruvate dehydrogenase E1 beta subunit | 4.846 | 0.401 | 5.247 |
| 34 | PDHA1 | pyruvate dehydrogenase E1 alpha 1 subunit | 4.406 | 0.407 | 4.813 |
| 35 | GOT2 | glutamic-oxaloacetic transaminase 2 | 1.275 | 0.042 | 1.317 |
| 36 | NDUFA9 | NADH:ubiquinone oxidoreductase subunit A9 | 1.025 | 0.259 | 1.284 |
| 37 | MDH1 | malate dehydrogenase 1 | 1.179 | 0.060 | 1.239 |
| 38 | NDUFA3 | NADH:ubiquinone oxidoreductase subunit A3 | 0.934 | 0.263 | 1.197 |
| 39 | NDUFA11 | NADH:ubiquinone oxidoreductase subunit A11 | 0.950 | 0.211 | 1.160 |
| 40 | NDUFS6 | NADH:ubiquinone oxidoreductase subunit S6 | 1.030 | 0.110 | 1.140 |
| 41 | MT-ND1 | mitochondrially encoded NADH:ubiquinone oxidoreductase core subunit 1 | 1.016 | 0.123 | 1.139 |
| 42 | NDUFS8 | NADH:ubiquinone oxidoreductase core subunit S8 | 1.007 | 0.109 | 1.116 |
| 43 | NDUFA5 | NADH:ubiquinone oxidoreductase subunit A5 | 0.982 | 0.122 | 1.104 |
| 44 | NDUFS2 | NADH:ubiquinone oxidoreductase core subunit S2 | 0.952 | 0.124 | 1.076 |
| 45 | NDUFB10 | NADH:ubiquinone oxidoreductase subunit B10 | 0.965 | 0.103 | 1.068 |
| 46 | NDUFA8 | NADH:ubiquinone oxidoreductase subunit A8 | 0.992 | 0.070 | 1.063 |
| 47 | NDUFB4 | NADH:ubiquinone oxidoreductase subunit B4 | 0.951 | 0.094 | 1.045 |
| 48 | NDUFB9 | NADH:ubiquinone oxidoreductase subunit B9 | 0.974 | 0.068 | 1.042 |
| 49 | NDUFC2 | NADH:ubiquinone oxidoreductase subunit C2 | 1.028 | 0.014 | 1.042 |
| 50 | NDUFA13 | NADH:ubiquinone oxidoreductase subunit A13 | 0.951 | 0.088 | 1.039 |
| 51 | MT-ND3 | mitochondrially encoded NADH:ubiquinone oxidoreductase core subunit 3 | 0.946 | 0.093 | 1.038 |

| Rank | Gene Symbol | Gene Name | Effect Size Lactate | Effect Size Succinate | Effect Size Sum |
|---|---|---|---|---|---|
| 52 | NDUFS1 | NADH:ubiquinone oxidoreductase core subunit S1 | 1.028 | 0.006 | 1.034 |
| 53 | NDUFA12 | NADH:ubiquinone oxidoreductase subunit A12 | 0.941 | 0.092 | 1.033 |
| 54 | NDUFA6 | NADH:ubiquinone oxidoreductase subunit A6 | 0.995 | 0.035 | 1.030 |
| 55 | NDUFC1 | NADH:ubiquinone oxidoreductase subunit C1 | 0.952 | 0.077 | 1.029 |
| 56 | NDUFB1 | NADH:ubiquinone oxidoreductase subunit B1 | 0.978 | 0.051 | 1.029 |
| 57 | NDUFB2 | NADH:ubiquinone oxidoreductase subunit B2 | 0.925 | 0.103 | 1.028 |
| 58 | NDUFA4 | NDUFA4 mitochondrial complex associated | 0.955 | 0.061 | 1.017 |
| 59 | NDUFA4L2 | NDUFA4 mitochondrial complex associated like 2 | 0.911 | 0.101 | 1.012 |
| 60 | NDUFS3 | NADH:ubiquinone oxidoreductase core subunit S3 | 0.965 | 0.045 | 1.010 |
| 61 | MT-ND4 | mitochondrially encoded NADH:ubiquinone oxidoreductase core subunit 4 | 0.941 | 0.061 | 1.002 |
| 62 | NDUFA2 | NADH:ubiquinone oxidoreductase subunit A2 | 0.968 | 0.019 | 0.988 |
| 63 | NDUFS7 | NADH:ubiquinone oxidoreductase core subunit S7 | 0.928 | 0.059 | 0.987 |
| 64 | MT-ND6 | mitochondrially encoded NADH:ubiquinone oxidoreductase core subunit 6 | 0.932 | 0.055 | 0.987 |
| 65 | NDUFA7 | NADH:ubiquinone oxidoreductase subunit A7 | 0.912 | 0.060 | 0.973 |
| 66 | NDUFAB1 | NADH:ubiquinone oxidoreductase subunit AB1 | 0.938 | 0.027 | 0.965 |
| 67 | NDUFA10 | NADH:ubiquinone oxidoreductase subunit A10 | 0.930 | 0.029 | 0.959 |
| 68 | MT-ND4L | mitochondrially encoded NADH:ubiquinone oxidoreductase core subunit 4L | 0.910 | 0.030 | 0.941 |
| 69 | NDUFB7 | NADH:ubiquinone oxidoreductase subunit B7 | 0.931 | 0.004 | 0.934 |
| 70 | MECR | mitochondrial trans-2-enoyl-CoA reductase | 0.072 | 0.172 | 0.244 |
| 71 | ACOT2 | acyl-CoA thioesterase 2 | 0.153 | 0.075 | 0.228 |
| 72 | FECH | ferrochelatase | 0.029 | 0.182 | 0.211 |
| 73 | MTHFD2L | methylenetetrahydrofolate dehydrogenase (NADP+ dependent) 2 like | 0.048 | 0.153 | 0.201 |
| 74 | GPD1 | glycerol-3-phosphate dehydrogenase 1 | 0.042 | 0.139 | 0.182 |
| 75 | NNT | nicotinamide nucleotide transhydrogenase | 0.056 | 0.125 | 0.181 |
| 76 | UCP2 | uncoupling protein 2 | 0.046 | 0.134 | 0.180 |
| 77 | ACO2 | aconitase 2 | 0.158 | 0.020 | 0.178 |
| 78 | ACSS1 | acyl-CoA synthetase short chain family member 1 | 0.165 | 0.012 | 0.176 |
| 79 | AADAT | aminoadipate aminotransferase | 0.019 | 0.132 | 0.150 |
| 80 | AMT | aminomethyltransferase | 0.010 | 0.138 | 0.148 |
| 81 | PPA2 | pyrophosphatase (inorganic) 2 | 0.073 | 0.073 | 0.146 |
| 82 | QDPR | quinoid dihydropteridine reductase | 0.010 | 0.135 | 0.145 |
| 83 | ALDH18A1 | aldehyde dehydrogenase 18 family member A1 | 0.070 | 0.068 | 0.138 |
| 84 | AOC1 | amine oxidase copper containing 1 | 0.044 | 0.079 | 0.122 |
| 85 | ODC1 | ornithine decarboxylase 1 | 0.068 | 0.050 | 0.118 |
| 86 | ATP8B2 | ATPase phospholipid transporting 8B2 | 0.031 | 0.087 | 0.118 |
| 87 | DHFR | dihydrofolate reductase | 0.031 | 0.080 | 0.111 |
| 88 | ATP11B | ATPase phospholipid transporting 11B (putative) | 0.085 | 0.026 | 0.110 |
| 89 | MPST | mercaptopyruvate sulfurtransferase | 0.017 | 0.092 | 0.109 |
| 90 | MCEE | methylmalonyl-CoA epimerase | 0.033 | 0.074 | 0.107 |
| 91 | GPD1L | glycerol-3-phosphate dehydrogenase 1 like | 0.020 | 0.084 | 0.104 |
| 92 | HAL | histidine ammonia-lyase | 0.063 | 0.032 | 0.095 |
| 93 | PTPMT1 | protein tyrosine phosphatase mitochondrial 1 | 0.007 | 0.080 | 0.087 |
| 94 | MAT2A | methionine adenosyltransferase 2A | 0.077 | 0.010 | 0.087 |
| 95 | HGD | homogentisate 1,2-dioxygenase | 0.026 | 0.058 | 0.084 |
| 96 | PGAM2 | phosphoglycerate mutase 2 | 0.035 | 0.038 | 0.073 |
| 97 | ADSSL1 | adenylosuccinate synthase like 1 | 0.016 | 0.054 | 0.069 |
| 98 | MCCC2 | methylcrotonoyl-CoA carboxylase 2 | 0.023 | 0.046 | 0.069 |
| 99 | AK3 | adenylate kinase 3 | 0.009 | 0.060 | 0.069 |
| 100 | MAOB | monoamine oxidase B | 0.020 | 0.034 | 0.054 |
| 101 | PCCA | propionyl-CoA carboxylase subunit alpha | 0.024 | 0.028 | 0.052 |
| 102 | UCP3 | uncoupling protein 3 | 0.032 | 0.011 | 0.043 |
